# The memory of pathogenic IgE is contained within CD23^+^IgG1^+^ memory B cells poised to switch to IgE in food allergy

**DOI:** 10.1101/2023.01.25.525506

**Authors:** Miyo Ota, Kenneth B. Hoehn, Takayuki Ota, Carlos J. Aranda, Sara Friedman, Weslley F. Braga, Alefiyah Malbari, Steven H. Kleinstein, Scott H. Sicherer, Maria A. Curotto de Lafaille

## Abstract

Food allergy is caused by allergen-specific IgE antibodies but little is known about the B cell memory of persistent IgE responses. Here we describe in human pediatric peanut allergy CD23^+^IgG1^+^ memory B cells arising in type 2 responses that contain peanut specific clones and generate IgE cells on activation. These ‘type2-marked’ IgG1^+^ memory B cells differentially express IL-4/IL-13 regulated genes *FCER2*/*CD23, IL4R*, and germline *IGHE* and carry highly mutated B cell receptors (BCRs). Further, high affinity memory B cells specific for the main peanut allergen Ara h 2 mapped to the population of ‘type2-marked’ IgG1^+^ memory B cells and included convergent BCRs across different individuals. Our findings indicate that CD23^+^IgG1^+^ memory B cells transcribing germline *IGHE* are a unique memory population containing precursors of pathogenic IgE.

**One-Sentence Summary:** We describe a unique population of IgG^+^ memory B cells poised to switch to IgE that contains high affinity allergen-specific clones in peanut allergy.

## Introduction

High affinity IgE antibodies are essential mediators of food allergy, a main cause of life-threatening anaphylaxis. Most food allergies develop in childhood and disproportionately affect children (*1*). A long-standing question in the allergy field is why allergies to some foods spontaneously resolve, while others persist. A key to understanding the evolution of food allergy may reside in the mechanisms that maintain the B cell memory of high affinity IgE responses. Murine studies demonstrated a lack of functional IgE memory cells and that high affinity pathogenic IgE plasma cells develop from the sequential switching of affinity matured IgG1^+^ memory B cells (*2, 3*). In humans, sequencing of the switch region of IgE^+^ B cells (*4*), the similarity of BCRs used by IgE^+^ and IgG^+^ B cells (*5, 6*), and longitudinal analysis of allergen-specific IgG and IgE antibodies (*7, 8*), also support a precursor role for IgG^+^ B cells in the generation of pathogenic IgE (*9*).

Peanut is the food allergen most often associated with anaphylaxis, and peanut allergy is generally persistent compared to other food allergies such as egg and cow’s milk allergy (*10-13*). Circulating levels of peanut specific IgE are predictive of thresholds of reactivity to peanut ingestion and whether peanut allergy will persist, though the relationship is imperfect, and other factors may play a role. In general, however, high peanut specific IgE levels (> 100 kU/L) are associated with long-term allergy persistence and reactivity to a small amount of the food, while low peanut specific IgE (< 5 kU/L) is associated with outgrowing peanut allergy and reactivity to a relatively higher amount of the food (*13-18*). We hypothesized that the existence of high affinity peanut specific IgG^+^ memory B cells and their ability to undergo class switching to IgE are critical for allergy persistence in pediatric peanut allergy.

Here we described in pediatric peanut allergy a population of IgG^+^ memory B cells that is marked by expression of IL-4/IL-13 regulated genes, contains high affinity allergen-specific clones, and is poised to switch to IgE.

## Results

### CD23^+^ IgG memory B cells frequency correlates with circulating IgE in pediatric peanut allergy

The expression of CD23, the low affinity receptor for IgE, was previously found to be increased in B lymphocytes of children with atopic dermatitis (AD) and asthma (*19, 20*), and the frequency of CD23^+^ B cells was found to correlate with IgE levels (*21*). To characterize B cells in food allergy, we first analyzed the expression of CD23 in B lymphocytes from PBMCs of 58 peanut allergic (PA) and 13 non-allergic (Non-PA) children. CD27 was used as a marker of memory B cells that are somatically mutated and likely to contain allergen-specific clones (*22*). Consistent with previous observations, we found a significantly higher percentage of CD23^+^ total, naïve, and memory B cells in PA than in non-allergic children of a similar age range (**Fig. 1A** and **Fig. S1A**). Possible correlations between CD23^+^ B cell populations and plasma IgE antibody concentration were analyzed in a cohort of PA children with a broad range of peanut specific IgE (**Figs. 1B-C, Fig. S1B**). CD27^+^ memory B cell populations were defined by expression of IgM/IgD (IgM/IgD^+^ unswitched, and IgM/IgD^-^ switched), and by IgG expression among IgM/IgD^-^ memory cells. A significant correlation was found between total IgE and CD23^+^IgM/IgD^+^ memory B cells, CD23^+^ IgM/IgD^-^ switched memory B cells, and CD23^+^IgG^+^ memory B cells (**Fig. 1B**). The levels of total IgE in circulation also correlated significantly with peanut specific IgE and with IgE specific for the main peanut allergen Ara h 2 (**Fig. S2A-B**). No correlation was found between the total IgE level and the frequency of CD23^+^ total B cells (**Fig. S2C**) or the CD23^-^ memory populations (**Fig. 1C**).

**Fig. 1.**
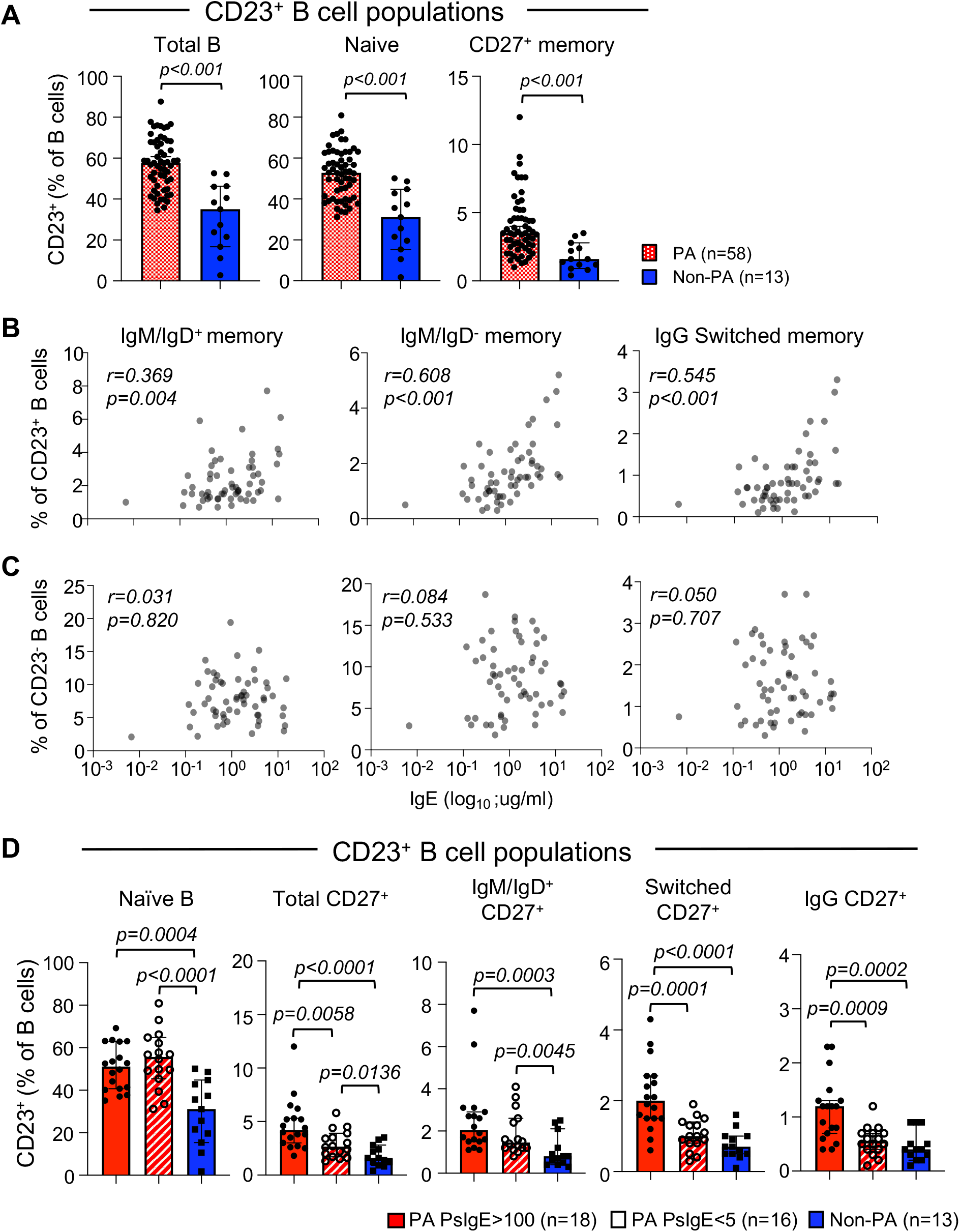
Frequency of IgG switched CD23^+^ memory B cells correlates with total IgE levels in PA. (**A**) Percentages of surface CD23^+^ B cells in total CD19^+^ B cells, in CD19^+^CD27^-^IgM^+^/IgD^+^ B cells (Naïve), and in CD19^+^CD27^+^ B cells (CD27^+^ memory) were analyzed in peanut allergic patients (PA, n=58; red dotted bar) and non-allergic healthy children (Non-PA, n=13; blue solid bar). *Mann-Whitney U test* was used for statistical analysis. (**B, C**) Frequency of switched CD23^+^ memory B cells showed a positive correlation with total plasma IgE levels while the frequency of CD23^-^ memory B cells did not show a significant correlation. *Spearman ranked correlation coefficient* was used for the statistical analysis. (**D**) CD23^+^ B cell populations were analyzed among peanut allergic patients with high specific IgE (PA PsIgE>100, n=18; red solid bar), with low specific IgE (PA PsIgE<5, n=16; red shaded bar) and Non-PA (n=13; blue solid bar). Statistical analysis was performed by *Mann-Whitney U test*.

A significant difference in total IgE levels was observed between children with a very high concentration of peanut specific IgE (PsIgE) in the blood (PsIgE > 100 kU/L referred to as PsIgE>100, n=18) and those with peanut specific IgE lower than 5 kU/L (referred to as PsIgE<5, n=16), while total IgE levels varied among PA children with intermediate peanut specific IgE (5<PsIgE<100) (**Fig. S1B**). To identify further differences in CD23 expression in the two ends of the spectrum of IgE sensitization, we compared the frequency of CD23^+^ B cells between PA children with PsIgE>100, PA children with PsIgE<5, and non-allergic children (Non-PA, n=13) (**Table S1**). The three groups had similar age ranges and median age (Methods). Both PA groups had a higher frequency of CD23^+^ naïve, CD23^+^ total memory, and CD23^+^ IgM/IgD^+^ memory B cells than the non-allergic groups. However, only the PsIgE>100 PA group had a significantly higher frequency of CD23^+^ total switched and CD23^+^ IgG^+^ memory B cells (**Fig. 1D** and **Fig. S2D**). Thus, although the frequency of CD23^+^ naïve B cells was similarly increased in PA children with very high or very low levels of peanut specific IgE, only highly sensitized children had a significantly higher frequency of CD23^+^ CD27^+^ total switched and IgG^+^ memory B cells.

### A cluster of somatically mutated switched memory cells express *FCER2, IL4R*, and *IGHE*

To further characterize B cell memory subsets, we performed 10X Genomics single-cell RNA sequencing (scRNA-seq) of whole transcriptome and B cell receptor (BCR) on flow-sorted CD27^+^ memory B cells from 5 PA children with PsIgE>100 and 3 non-allergic children (**Table S2**). We identified 10 clusters of memory B cells (numbered 0 to 9) and identified differentially expressed genes (DEG) in each cluster (**Fig. 2A-B, Figs. S3-4**). Although genes for antibody isotypes except *IGHE* were not used in clustering, BCR analysis identified clusters enriched in unswitched IgM memory B cells (clusters 3 and 9), switched memory cells (clusters 1, 4, 5, and 8), or containing a mixture of isotypes (clusters 0, 6, and 7) (**Fig. 2C, Fig. S5**). Clusters 0 and 2 share main DEGs such as *YPEL, NR4A2, H3F3B*, and *FOSB. CD83* expression marked clusters 2, 6, and 9, and other main DEGs of cluster 2 included *NR4A1, FOS*, and *JUNB*. Cluster 3 DEGs included *FGR* and *CD1C*, cluster 8 DEGs included *BTG2*, and other DEGs of clusters 7 and 8 included mainly ribosomal proteins and RNA binding proteins. The small cluster 9, which is highly enriched for *IGHM* BCRs, was characterized by high expression of *EGR1/2/3* and *MYC*, suggesting recent activation. Using mass cytometry Glass et al. characterized memory populations by expression of CD27, CD45RB, CD73, CD95, or CD11c (*23*). Our transcriptional analysis was done on CD27^+^ memory B cells, and it did not distinguish *CD45* isoforms. We detected *CD73/NT5E* expression in only a small percentage of cells in clusters 4, 5, and 6, and transcripts for *FAS/CD95* and *CD11c/ITGAX* were undetectable in most cells (**Fig. S6**). Cluster 3 expressed some marker genes of atypical memory B cells (*24*), such as *FGR, and FCRL5*, and had low expression of *CR2/CD21* (**Fig. S6**).

**Fig. 2.**
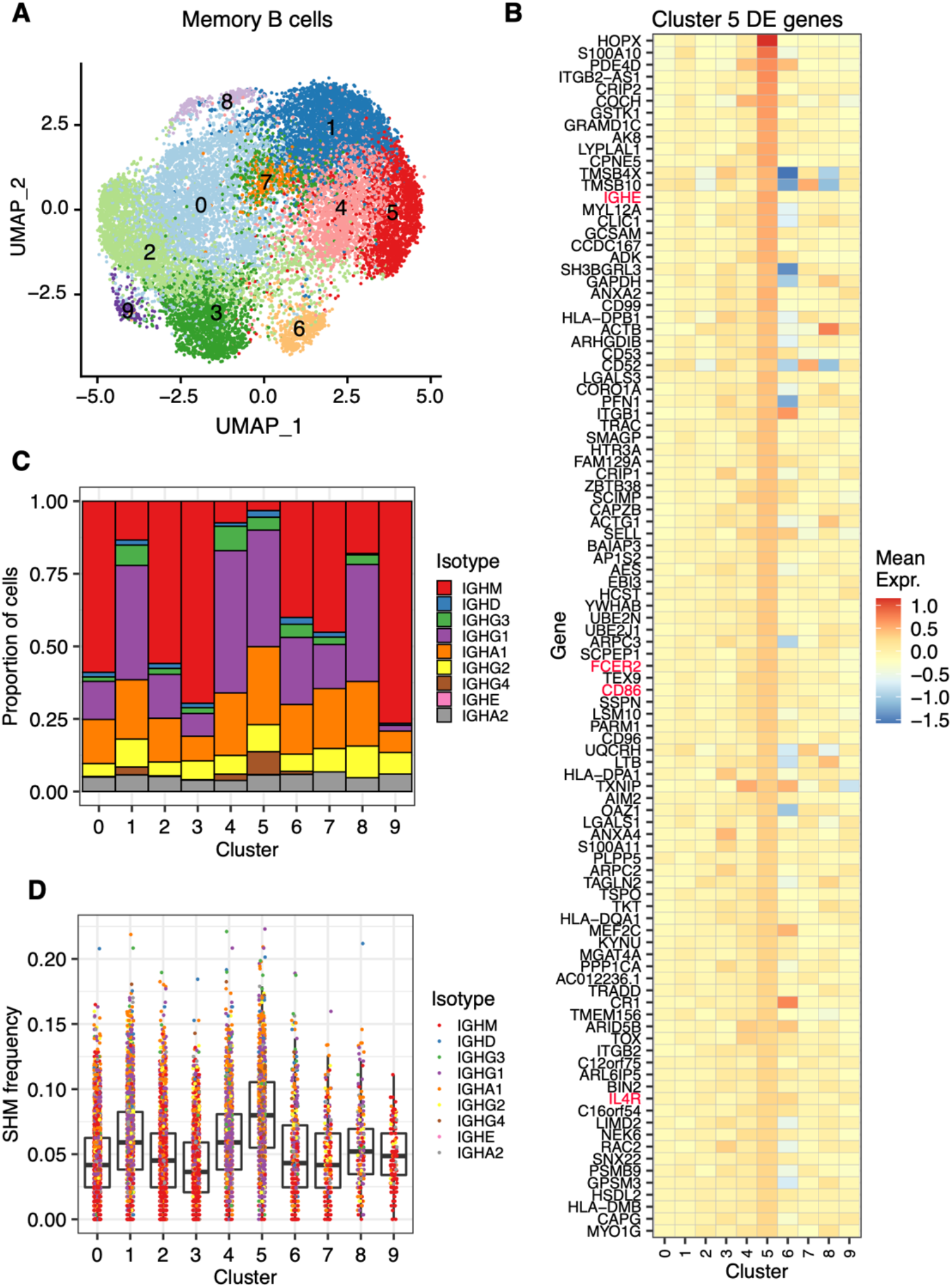
scRNA-seq characterization of CD27^+^ memory B cells in PA and non-PA subjects. (**A**) UMAP of cells obtained from peanut allergic (PA) and healthy control (non-PA) subjects. Each point represents a single cell. (**B**) Scaled gene expression of cells in cluster 5. Tiles show scaled, normalized expression of cluster 5. Genes are positively differentially expressed genes (adjusted p < 0.05) for cluster 5. (**C**) Proportion of cells in each scRNA-seq defined cluster that uses each isotype constant region. Cluster 5 was highly enriched for *IGHG*-associated BCRs (61.7% of cells, *p < 2*.*2e-16 chi-squared test*). (**D**) Level of IGHV-gene SHM for cells in each cluster. Cells are colored by associated constant region.

Given the close association between CD23^+^ IgG^+^ memory B cells and peanut allergy (**Fig. 1**), we searched for a cluster that contained CD23^+^ IgG^+^ memory B cells. Cluster 5 was marked by high levels of *FCER2* (the gene that encodes CD23) (**Fig. 2B, Fig. S6**) and contained mostly IgG and IgA cells (**Fig. 2C)**. *FCER2* is upregulated by IL-4/IL-13 signaling in B cells, and importantly, clusters 5 DEGs included other IL-4/IL-13 regulated genes such *IL4R, CD86* and *IGHE*. Thus cluster 5 contains type2-marked memory cells similar to the ones we recently described (*25*). Top DEGs of cluster 5 included *HOPX* (**Fig. 2B**), encoding a homeobox-domain protein previously found to be associated with IgG^+^ memory B cells and naïve B cells (*26*), *S100A10*, encoding a calcium-binding Annexin A2 partner protein that regulates the activity of many transmembrane proteins (*27*), *ANXA2* (encoding Annexin A2) and genes involved in MHC II antigen processing/presentation *HLA-DPB1, HLA-DPA1, HLA-DQA1, HLA-DMB, and SCIMP* (**Fig. 2B**). Cluster 5 DEGs shared with clusters 4 and 6 were *PDE4D*, encoding phosphodiesterase 4D, *TXNIP*, encoding thioredoxin interacting protein, *SELL*, encoding selectin B and *ARID5B*, encoding a transcription regulator of B lymphocytes. Cluster 5 DEGs shared with cluster 4 were *COCH*, encoding cochlin, and *TOX*, a transcription factor expressed by germinal center cells and CD80^+^CD73^+^ IgG1 mouse memory cells (*3*). DEGs in the type2-marked memory cluster recently identified (*25*) included *FCER2, IL4R, IL13RA1, RNGTT, HOPX, CD74, PARM1* and *SCIMP*. Cluster 5, and to a lesser extent clusters 4 and 6, expressed these genes, indicating it is likely a corresponding population to the type2-marked memory cluster in our data (**Fig. S6)**.

To further understand the biological processes shaping the memory B cells in cluster 5, we performed gene ontology enrichment analysis. The most-enriched gene ontologies for cluster 5 were related to the cytokine-mediated signaling pathway (GO: 009221), and antigen receptor-mediated signaling pathway (GO: 0050851) (**Table S3**). We next determined whether memory B cells in cluster 5 from PA subjects had different expression signatures than cluster 5 cells from non-PA subjects. We found that PA cells in cluster 5 were enriched for cytokine-mediated signaling pathway signatures (GO: 009221) (**Table S4**). Overlap with this gene ontology included genes involved in signal transduction through JAK-STAT and JAK-IRS2 (*JAK1, PIM1, JUNB, CDKN1B, GRB2, HNRNPF, PTPN6*, and *SOCS1*). Thus cluster 5 in PA subjects was enriched for transcriptional signatures consistent with cytokine signaling, MHC II receptor activity, and JAK signaling.

We further characterized cluster 5 using BCR repertoire sequencing. Cluster 5 was highly enriched for *IGHG*-associated BCRs (61.7% of cells), predominantly *IGHG1* (40.1% of cells), as well as the highest proportion of *IGHG4*-associated BCRs among all clusters (7.84%) (**Fig. 2C**). BCRs in cluster 5 were highly mutated, with average SHM frequency of 8.08%, the highest of any cluster (**Fig. 2D**). This was not due to the low frequency of IGHM cells: *IGHG1, IGHG2*, and *IGHG3* cells in cluster 5 were more mutated than their respective cell types in other clusters (**Fig. S7**). While true IgE cells are rare, we identified two B cells with *IGHE* BCRs, each from a different PA subject and both belonging to cluster 5 (**Fig. S7**). Further, both had detectable *FCER2* expression and high levels of SHM (7.3% and 11.1%).

In addition to the immunoglobulin constant region associated with a BCR, B cells can produce non-coding transcripts from the constant immunoglobulin genes. These could be germline transcripts indicating priming for near-future class switching between isotypes, or a past rearrangement in the non-productive *IGH* allele (*28-30*). In the scRNAseq experiment, we identified 178 B cells with normalized *IGHE* transcript levels at least as high as the two B cells found with *IGHE* BCRs (**Fig. 3A**). These *IGHE* transcribing B cells had non-*IGHE* constant regions associated with their BCRs, predominantly *IGHG1* or *IGHG4* (62.4% and 18.5% of *IGHE* transcribing cells, respectively) and were primarily found in cluster 5 (57.3 % of *IGHE* transcribing cells, **Fig. 3B**), suggesting that cluster 5 contains IgG^+^ memory B cells primed to switch to IgE.

**Fig. 3.**
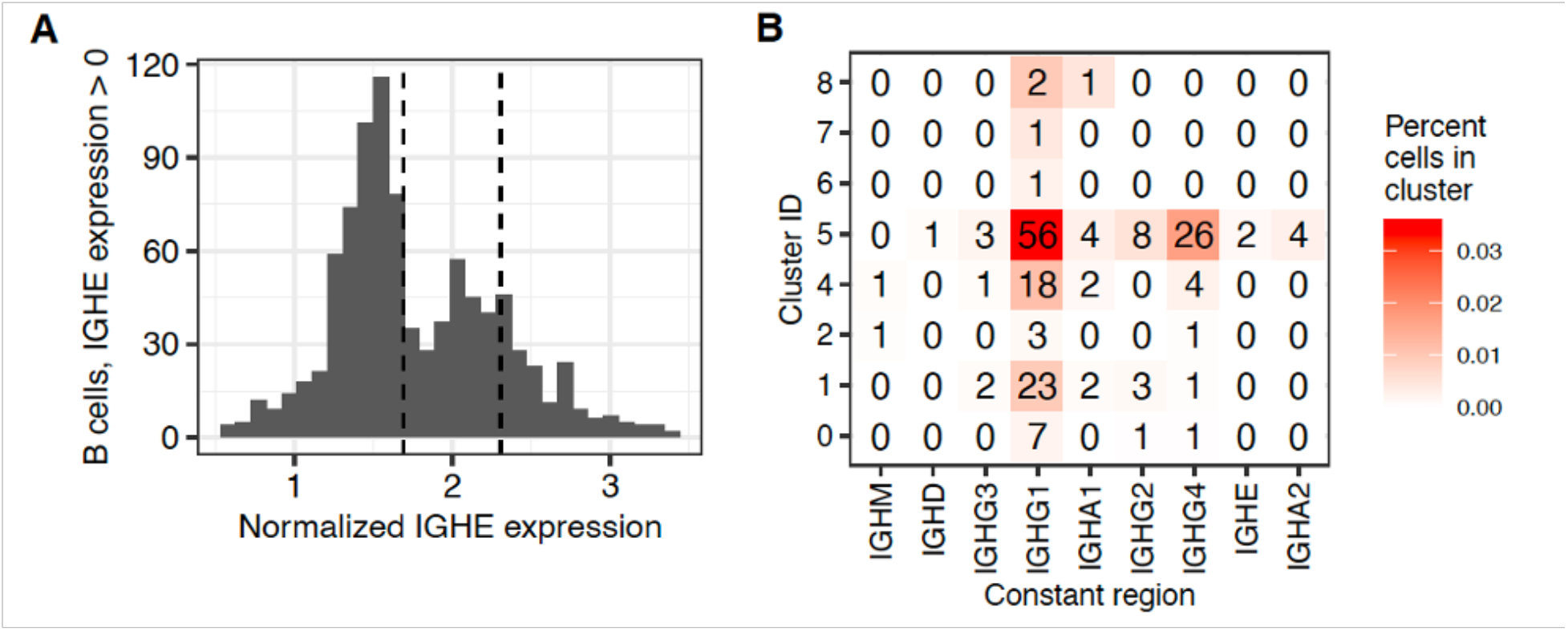
*IGHE* transcribing B cells are associated with cluster 5. **(A)** Distribution of normalized IGHE expression among all B cells that express at least one *IGHE* transcript. Dashed lines show the level of *IGHE* transcription in the two observed *IGHE*-constant region B cells. (**B**) *IGHE*-transcribing cells associated with each cluster and constant region. Cells of the grid are labeled by the total number of *IGHE*-transcribing cells and colored by the percentage of cells in each cluster that are *IGHE*-transcribing. *IGHE* transcribing B cells used *IGHG1* or *IGHG4* constant regions in their BCRs were primarily found in cluster 5 (57.8% of *IGHE* transcribing cells, *p < 2*.*2e-16, by chi-squared test*).

### Peanut specific B cells in IgG memory populations

We subsequently asked if CD23^+^IgG^+^ memory B cells contained peanut specific clones, and whether their frequency was higher in PBMC from highly sensitized PA children (PsIgE>100) than in those with low peanut specific sensitization (PsIgE<5). Since *IL4R* expression was transcriptionally upregulated in cluster 5 (**Fig. 2B**), which contained *FCER2/CD23* expressing IgG memory cells, we defined a sorting gate that comprised all cells with detectable expression of CD23 and IL4R and referred to this population as CD23^+^/IL4R^+^. CD23^+^/IL4R^+^IgG^+^ memory B cells and CD23^-^IL4R^-^IgG^+^ memory B cells were sorted and cultured at 1,000 cells per well over fibroblasts expressing CD40L and BAFF with the addition of the cytokines IL-2, IL-4, IL-10 and IL-21 **(Fig. 4A and Fig. S8A**). These conditions were optimized for B cell proliferation and plasma cell differentiation which also allowed class switching to IgE. After 7 days, peanut specific antibodies were measured in the culture supernatants, and cells were analyzed by flow cytometry. Most wells of cells from non-allergic subjects were negative for peanut specific IgG antibodies (**Fig. 4B**, right). In cultures from PA children with low specific IgE (PsIgE<5), the frequency of positive wells for peanut specific IgG were similar between CD23^-^IL4R^-^ and CD23^+^/IL4R^+^IgG^+^ cell cultures (**Fig. 4B**, middle). Importantly, in cultures of cells of PA children with high specific IgE (PsIgE>100), wells containing peanut specific IgG antibodies were predominantly found among CD23^+^/IL4R^+^IgG^+^ memory cell cultures (**Fig. 4B**, left). In addition, IgG^+^ memory B cells from PA children with high specific IgE (PsIgE>100) proliferated more than cells from the other two groups (**Fig. S8B**). Class switching to IgE was relatively low in these culture conditions, nevertheless, we detected the highest frequency of IgE B cells in the cultures of CD23^+^/IL4R^+^ IgG memory B cells from PA children with high specific IgE (PsIgE>100) (**Fig. S8C**). Thus, in highly sensitized peanut allergic children, peanut specific IgG memory clones are enriched in a CD23^+^ population with the ability to switch to IgE.

**Fig. 4.**
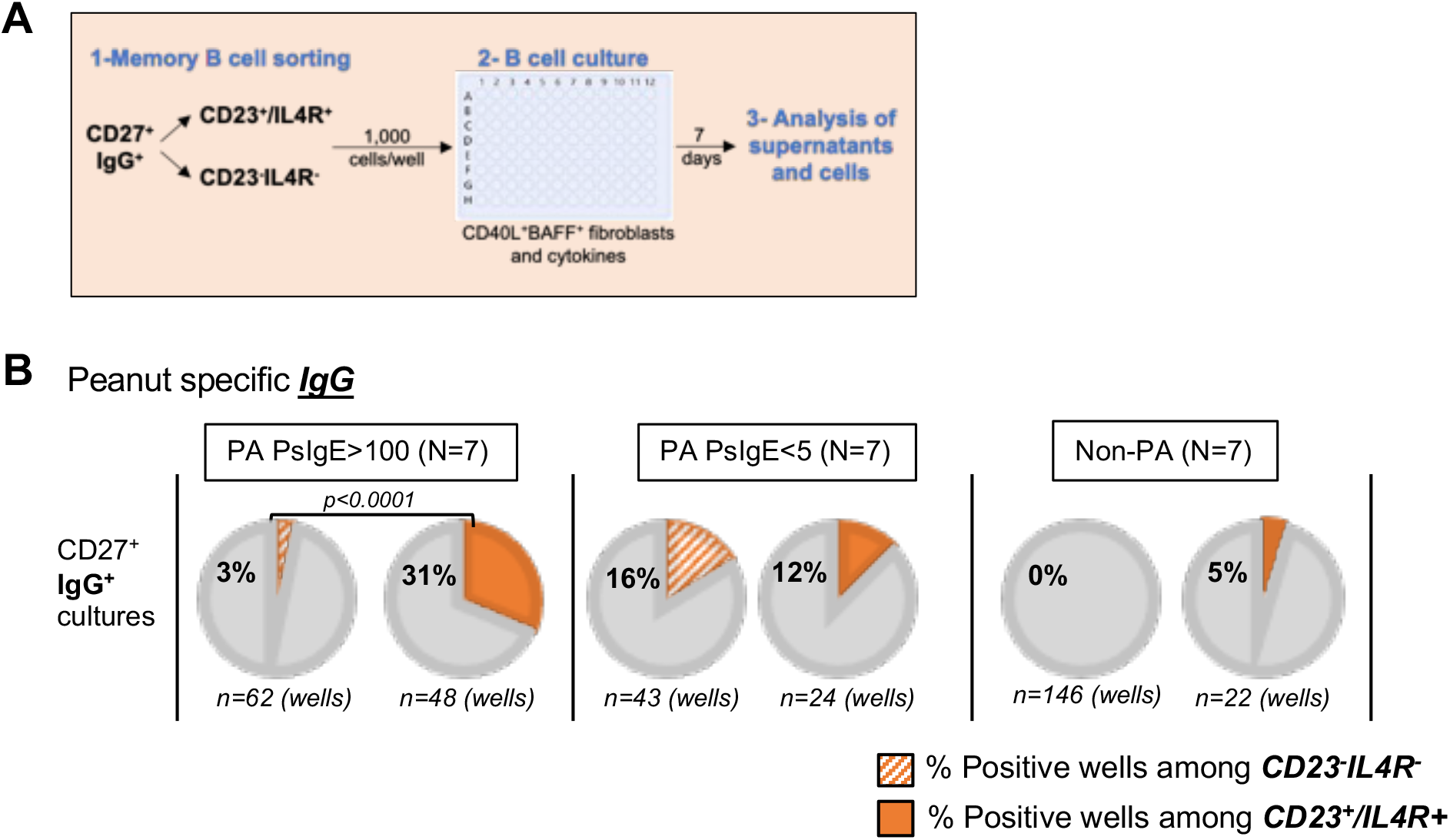
IgG switched CD23^+^ memory B cells from PA with sIgE>100 showed an increased frequency of B cells producing peanut specific IgG. **(A)** Two distinct populations of CD27^+^ memory B cells were sorted for memory B cell culture experiments. **(B)** Culture supernatants of IgG^+^ memory B cells from PA with PsIgE>100 (N=7, left), PA with PsIgE<5 (N=7, middle), and Non-PA (N=7, right) were tested for the reactivity against peanut by ELISA. Percent of positive wells among the total wells (indicated in each circle) of each culture is shown. Shaded orange pie represents a percent positive among CD23^-^IL4R^-^ memory B cell culture wells and solid orange pie represents a percent positive among CD23^+^ memory B cell culture wells. PA with PsIgE>100 showed a higher percentage of positive wells in cultures of CD23^+^ than in CD23^-^IL4R^-^IgG^+^ memory B cells (*p=0*.*00006* by *Chi-square test*).

### Ara h 2 sorted IgG1 B cells are highly mutated and transcribe *FCER2* and germline *IGHE*

To determine at the single cell level if peanut specific memory B cells were contained within an IgG population with the characteristics of the cells described in cluster 5, we isolated from PA patients B cells that bound the main peanut allergen Ara h 2 (*31-33*). Ara h 2 binding B cells were single-cell sorted from PBMC of 8 PA subjects with PsIgE>100 and 5 PA subjects with PsIgE<5 using fluorescent Ara h 2 multimers. For comparison, we sorted diphtheria toxin (DT) binding B cells from PA and non-allergic children (Non-PA), as these cells are expected to be found in most children due to widespread vaccination (**Table S5, Fig. S9A)**. The frequency of Ara h 2 binding B cells was significantly higher in PA with PsIgE>100 than in PA with PsIgE<5, while Ara h 2 binders were undetectable in non-allergic samples. DT binders were detected at similar frequencies among the three groups (**Fig. S9B**).

To characterize the BCRs of Ara h 2 binding B cells, heavy and light chain BCR genes were sequenced from cDNA synthesized from sorted single cells. In total, we obtained heavy chain sequences from 69 Ara h 2-binding B cells and 174 DT-binding B cells, 86.4% of which had at least one associated light chain sequence. Heavy chain sequences of Ara h 2-binding IgG B cells from PA subjects were significantly more mutated than DT-binding IgG B cells from both PA and non-PA subjects (**Fig. 5A**). By contrast, most Ara h 2- and DT-binding IgM cells were unmutated (**Fig. 5A**). Furthermore, Ara h 2-binding IgG B cells from PA subjects were more likely to be of the IgG1 sub-isotype (85.7% of *IGHG* cells, **Fig. 5B**), compared to DT-binding IgG1 B cells from PA (55.1% of *IGHG* cells) or from non-allergic subjects (52% of IGHG cells).

**Fig. 5.**
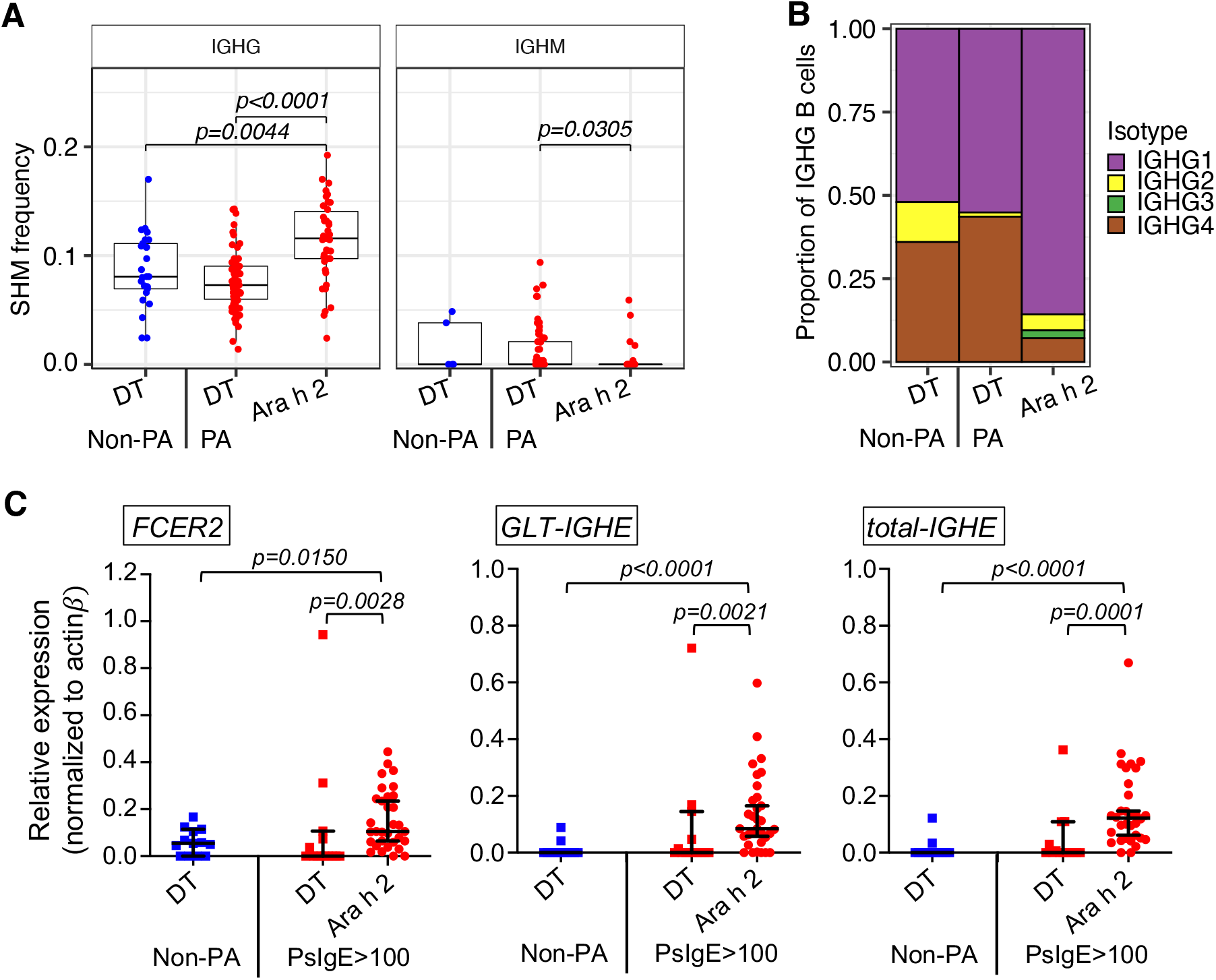
BCRs from Ara h 2-binding IgG1^+^ B cells are highly mutated, use *IGHG1* constant regions, and express *FCER2* and *IGHE*. (**A**) Level of IGHV-gene somatic hypermutation for DT and Ara h2-sorted B cells with either *IGHG* or *IGHM* constant regions from either PA or non-PA subjects. Ara h 2 binding IgG^+^ B cells were more mutated compared to DT binging IgG^+^ B cells from PA and non-PA (*Mann-Whitney U test*). (**B**) Proportion of DT and Ara h 2-sorted *IGHG* B cells with associated *IGHG* sub-isotypes. Ara h 2 binding B cells from PA were more likely to be IgG1 compared to DT-binding B cells from PA (85.7% vs 55.1% of IGHG, *p = 0*.*0015*) and non-PA (85.7% vs 52% of IGHG, *p=0*.*006 by chi-square test*). (**C**) *FCER2, GLT-IGHE*, and total-*IGHE* gene expression levels in single sorted IgG1^+^ B cells. mRNA was quantified in single IgG1^+^ B cells derived from Ara h 2 binders (n=34 cells) from 6 PA subjects with PsIgE>100 and DT binders from 6 PA subjects with PsIgE>100 (n=14 cells) and 3 non-PA (n=23 cells). Ara h 2 binding IgG1^+^ B cells showed higher *FCER2, GLT-IGHE*, as well as *total-IGHE* expression compared to DT binding IgG1^+^ B cells from PA and non-PA (*Mann-Whitney U test*).

To determine if Ara h 2-binding IgG1 cells belonged to the CD23^+^IgG^+^ memory population characterized through scRNAseq (cluster 5; **Fig. 2**), we used real-time PCR to quantify *FCER2/CD23*, germline *IGHE* (*GLT-IGHE*) and total-*IGHE* transcripts in all antigen-binding IgG1cells from PA subjects with PsIgE>100 and from non-allergic subjects (**Fig. 5C**). *FCER2/CD23* expression was detected in most Ara h 2-binding IgG1 cells from PA subjects, and its average expression was significantly higher in Ara h 2-binding IgG1 cells than DT-binding IgG1 cells from PA or non-allergic subjects. Importantly, total *IGHE* and germline *IGHE* transcripts were detected in most Ara h 2-binding cells but few DT-binding cells. These results show that Ara h 2 binding IgG B cells from PA subjects have defining characteristics of type-2 marked IgG cells from cluster 5: high somatic hypermutation, predominant IgG1 constant regions, and transcription of *FCER2/CD23* as well as *IGHE*. The fact that Ara h 2 binding IgG1 memory cells transcribe germline *IGHE* is particularly relevant, as germline transcription is an essential initial step for class switch recombination (*29*).

### Identification of high affinity convergent Ara h 2 specific BCR sequences across different subjects

BCRs within individual humans are highly diverse, and identification of similar sequences among different individuals may be indicative of convergent antigen-driven selection. To identify convergent BCRs specific for Ara h 2, we performed clustering based on IGH amino acid sequence similarity (Methods) using the BCR sequencing data from Ara h 2- and DT-binders, and from all memory B cells profiled in the 10X scRNA-seq experiment. We refer to B cells with highly similar BCRs in multiple subjects as “convergent sequence families.”

Among cells isolated by Ara h 2 binding, monoclonal antibodies derived from a convergent sequence family of 2 IgM memory cells (P5AC4, P5EC1) from two PA subjects, as well as an antibody derived from an unrelated IgM Ara h 2 binder (P2CC6), displayed very low binding to Ara h 2 in ELISA. Of these, only P2CC6 had some very low reactivity against Ara h 1 and Ara h 3 (**Fig. S10A**). The heavy chains of these three IgM cells had low SHM frequencies (between 0 and 2.1%).

Among Ara h 2 binding IgG cells, we identified a group of convergent sequences composed of three IgG1 B cells (P4EC10, P4AC5, and P6EC2) using ambiguously IGHV3-30*18 or IGHV3-30-5*01 paired with IGHJ6*02, as well as IGKV3-20*01 and either IGKJ2*01 or IGKJ2*03. These cells were from three different PA subjects and share between 73.4 and 78.2% amino acid identity with a previously described (PA13P1E10) anti-Ara h 2 IgE plasma cell (*34*) (**Fig. 6A**, PA13P1E10 family). Further, all three heavy chains were highly mutated, with SHM frequencies between 9.7% and 14.9%. To confirm whether the cells from this convergent group of sequences bound peanut antigens, we generated recombinant human IgG1 antibodies. In ELISA and biolayer interferometry assays, we found that these antibodies bound with high affinity to Ara h 2 (KD (M): <1.0E-12 for P4EC10) and cross-reacted with Ara h 1 and Ara h 3 (**Fig. 6B and Fig. S10B, S11A-B)**.

**Fig. 6.**
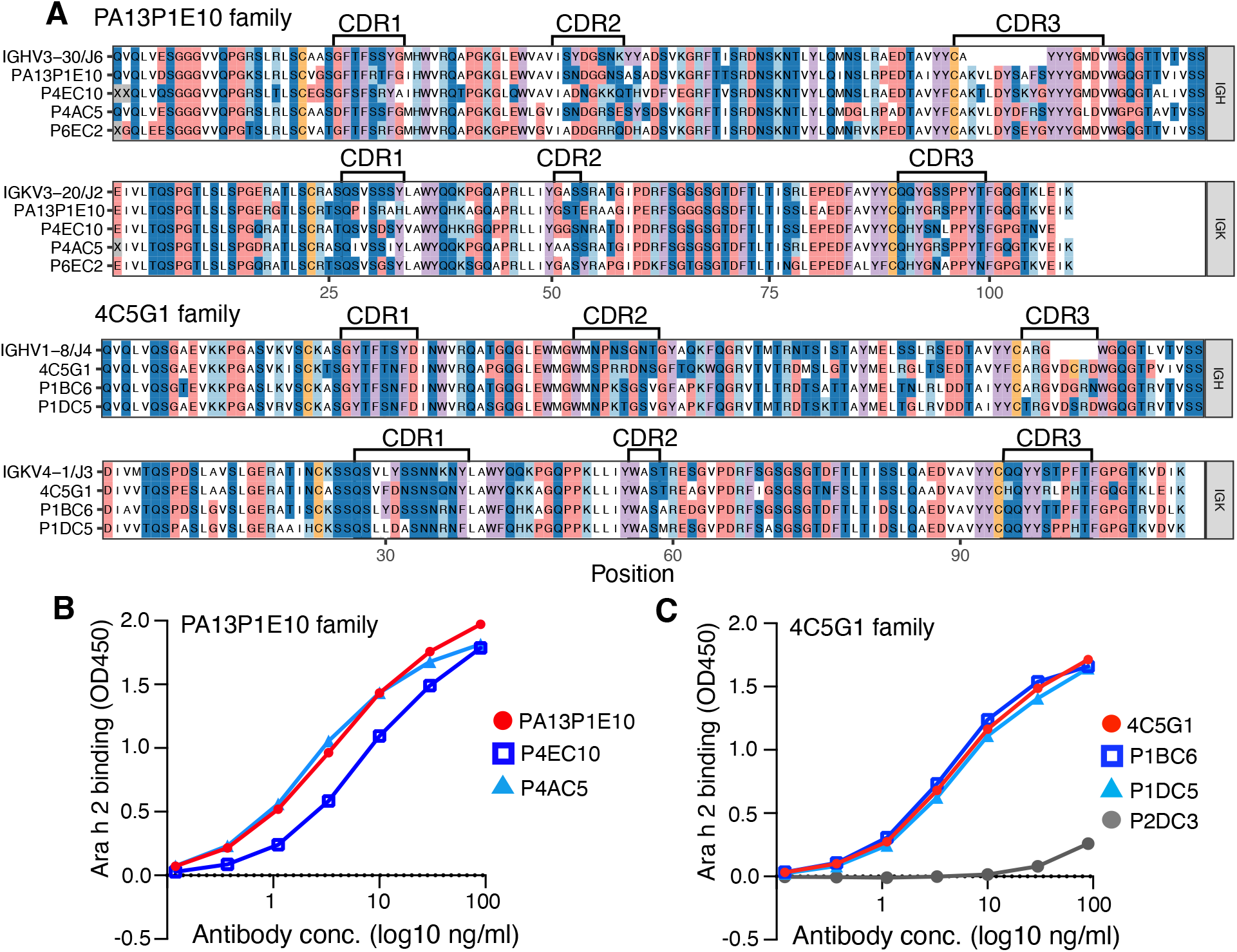
Convergent Ara h 2-binding sequence cluster across multiple donors. (**A**) Amino acid alignment of heavy and light chain sequences from two shared convergent sequence clusters. The upper sequence in each plot shows the predicted germline V and J gene sequences, followed by clonal members. Brackets indicate CDRs on sequence alignments. (**B-C**) ELISA curve for monoclonal antibodies generated from V(D)J sequences of convergent clusters from panel **A**.

A second convergent family of Ara h 2-binding IgG sequences mapped back to cluster 5 in the scRNAseq data (**Figs. 2-3**). It contained three cells from two PA subjects: one cell from cluster 5 of the 10X scRNAseq (4C5G1) and two sorted Ara h 2 binder cells (P1BC6 and P1DC5, **Fig. 6A**, 4C5G1 family). Importantly, cell 4C5G1 from scRNAseq cluster 5 had detectable transcripts of both *FCER2* and *IL4R*. All three cells in this cluster were used with IGHV1-8*01 and IGHJ4*02, as well as IGKV4-1*01 and either IGKJ3*02 or IGKJ2*01. These cells shared 73.0 to 88.6% amino acid identity in the heavy chain. Furthermore, all three heavy chain sequences had high SHM levels (between 11.5% and 14.2%) and all three bound Ara h 2 with high affinity (KD (M): 3.42E-11 for 4C5G1; 1.13E-9 for P1BC6; 9.09E-11 for P1DC5) and cross-reacted with peanut antigens Ara h 1 and Ara h 3 (**Fig. 6C** and **Fig. S10C, S11A-B**).

These results indicate that in highly sensitized PA subjects, high affinity B cell clones for the main allergen Ara h 2, are predominantly somatically mutated IgG1 memory cells that transcribe germline *IGHE* and contain convergent BCR specificities across different subjects.

## Discussion

Using single-cell transcriptomics and functional analysis we identified a population of somatically mutated IgG memory cells characterized by expression of IL-4/IL-13 regulated genes *CD23/FCER2, IL4R*, and *IGHE*, composed mostly of IgG1 and IgG4 cells. This IgG memory population is very similar to the type2-marked IgG memory B cells that we recently described and showed were increased in adult subjects with atopic disease (*25*), and to a reported small population of double negative B cells (*35*). Expression of *IL4R* suggests a readiness to respond to IL-4, and transcription of *IGHE* is particularly striking as germline *IGHE* transcription is essential to initiate class switch recombination to IgE (*29*). The frequency of CD23^+^IgG^+^ memory B cells in PBMC correlated with circulating IgE levels in a peanut allergic pediatric population, which also supports a relationship between these memory cells and IgE. In fact, IgG memory cells expressing CD23, obtained from highly sensitized PA children, contained peanut specific clones and gave rise to IgE cells in vitro. Furthermore, IgG^+^ memory B cells that bound the main peanut allergen Ara h 2 with high affinity were predominantly IgG1 cells expressing *FCER2* and germline *IGHE*, thus belonging to the *FCER2*^*+*^ *IGHE*^+^ IgG population identified in cluster 5 of the 10X scRNAseq analysis. In line with our model of IgE cell differentiation developed from mouse studies (*2, 3, 9, 36*), we proposed that allergen-specific IgG^+^ memory B cells expressing *CD23*/*FCER2* and germline *IGHE* are precursors of pathogenic IgE plasma cells in highly sensitized peanut allergic subjects.

Pathway analysis of differentially expressed genes of the *IGHE*^+^ memory cluster (cluster 5) demonstrated enhanced expression of genes related to cytokine-mediated signaling and antigen receptor mediated signaling. Within cluster 5, the cytokine-mediated signaling pathway was increased in PA samples compared to non-allergic samples, and in particular, the expression of *JAK1, GRB2, PTPN6*, and *SOCS1* was higher in PA samples. This suggests that differential activation in PA cells of IL4R-mediated JAK1-STAT6 signaling, which promotes class switching to IgE (*29*), as well as JAK1-IRS2 signaling, which induces survival and proliferation of B cells via Grb2-mediated activation of PI3K and ERK1/2 (*37, 38*). The increased expression of inhibitors *PTPN6* and *SOCS1* also supports the higher activity of the IL4R pathway, as JAK1-STAT6 activation is controlled by SHP-1 (*PTPN6*) (*39*), and the JAK1-IRS2 pathway is controlled by *SOCS1* (*40*). Compatible with the DEG analysis of cluster 5, CD23^+^/IL4R^+^IgG^+^ memory B cells from highly sensitized peanut allergic children had high proliferation and generated the most IgE cells after 7 days of culture compared with other samples, suggesting enhanced IL4R-JAK1 signaling. JAK inhibition may thus directly suppress type2-marked IgG^+^ memory B cell activation and class switching to IgE. Importantly, a JAK inhibitor that is FDA approved for the treatment of atopic dermatitis (*41*) is now being tested for the treatment of food allergy (clinical trial NCT05069831).

Several MHC II genes and genes related to the MHC II pathway were increased in the *IGHE*^+^ cluster 5, suggesting that those cells had high competence for antigen presentation to CD4 T cells. Other DEGs with high significance were *HOPX* and *S100A10. HOPX* encodes an atypical homeobox protein that binds serum response factor (SRF) (*42*). HOPX function in B cells is not known but it was reported to be expressed in IgG^+^CD27^+^ B cells in the spleen (*26*) and in pre-plasmablasts of human tonsils (*43*), thus *HOPX* may mark high affinity switched memory cells prone to the plasma cell fate.

Increased CD23 expression in B lymphocytes has been associated with allergic diseases and correlated with IgE production (*19-21*). It has been shown that IgE stabilizes the trimer of membrane CD23 on the cell surface and protects it from proteolysis (*44*). Thus, in addition to increased transcription, stabilization of CD23 by IgE in allergic patients may contribute to higher membrane expression. Interestingly, circulating IgE plasmablasts had higher expression of MHC II genes and *FCER2* than plasmablasts of other isotypes (*34*). While this was considered evidence of lower maturation of IgE cells, it could reflect a developmental relationship between IgE cells and their potential precursors, CD23^+^IgG1^+^ memory B cells.

Previous repertoire analysis of circulating B cells found that the IgE repertoire was most closely related to IgG1, suggesting sequential class switching from IgG1 to IgE (*45*). In subjects undergoing SLIT pollen immunotherapy, shared clonotypes between IgE and IgG BCR sequences strongly indicated common clonal origin (*6*). A recent study described repertoire relatedness between IgA1 and IgE in gastrointestinal tissue from peanut allergic patients, suggesting that IgA1 to IgE switching could take place in the gut (*46*). However, in our 10X scRNAseq analysis, very few IgA1^+^ or IgM^+^ cells expressed *IGHE*, and *IGHE* transcription was strongly associated only with IgG1^+^ and IgG4^+^ memory B cells. It is also worth noting that antibodies derived from Ara h 2 binding IgM clones showed low or no somatic hypermutation and very low binding to Ara h 2. Ara h 2-specific IgE has been reported to correlate with disease severity in peanut allergy (*47*). We found that the highly sensitized PA allergic children had a much higher frequency of Ara h 2-binding IgG1^+^ memory B cells than children with low levels of peanut specific IgE, again suggesting a relationship between peanut specific IgG1 memory cells and specific IgE antibodies. Consistent with our results, other studies also found relatively high mutation rates in Ara h 2-specific antibody genes (*34, 48, 49*), and identified convergent sequence families of Ara h 2 binding BCRs, suggesting preferential binding of the Ara h 2 protein to selected germline VH and VL specificities that undergo further affinity maturation across individuals.

In sum, we describe a population of somatically mutated CD23^+^*IGHE*^*+*^IgG1^+^ memory B cells with a high potential to respond to IL-4 and to undergo class switching to IgE. This population contains high affinity Ara h 2 specific memory B cells in highly sensitized peanut allergic individuals. We propose that CD23^+^*IGHE*^+^IgG1^+^ memory B cells are involved in the persistence of food allergy by providing precursors of pathogenic IgE plasma cells. Thus, CD23^+^*IGHE*^+^IgG^+^ memory B cells are a potential therapeutic target in peanut allergy.

## Acknowledgments

We thank Jiaming Lin and clinical personnel of the Jaffe Food Allergy Institute and General Pediatrics at ISMMS for patient recruitment; Ankita Prakash for general lab support; the Human Immuno Monitoring Center, and the Genomics and Flow Cytometry Cores at ISMMS for technical support; James Duty for help with Octet experiments and Juan Lafaille for critical comments. M.O. received a fellowship from the ISMMS T32 National Institute of Health (NIH) training program. This work was funded by the National Institute of Allergy and Infection Diseases grants R01AI104739 (S.H.K), K99AI159302 (K.B.H.), R01AI151707 (M.A.C.L.) and R01 AI153708 (M.A.C.L.)

## Author contributions

M.O. designed and carried out most of the experiments in this manuscript in consultation with M.A.C.L. K.B.H. performed all computational analysis for whole transcriptome expression and BCR repertoire with S.H.K. supervision. T.O. performed BCR repertoire and mutation analysis of cultured B cells and contributed to critical discussions. S.F. performed antibody binding experiments. W.F.B. performed single cell expression analysis. S.H.S. and A.M. provided overall clinical oversight for allergic (S.H.S.) and non-allergic (A.M.) subject recruitment. C.J.A. contributed to critical discussions. M.O., K.B.H., and M.A.C.L. wrote the manuscript. M.A.C.L. supervised the project. All authors edited and approved the manuscript.

## Data and materials availability

Scripts used to perform analysis are available at https://bitbucket.org/kleinstein/projects.

Single-cell RNA sequencing data of this manuscript is contained in BioProject PRJNA847159, and GEO accession GSE208235.

## Supplementary Materials

Methods

Figs. S1 to S11

Tables S1 to S7

Data S1

References (*50–62*)

