## Supplementary Materials and Figures for "The memory of pathogenic IgE is contained within CD23^+^IgG1^+^ memory B cells poised to switch to IgE in food allergy"

**This PDF file includes:**

Methods

Figs. S1 to S11

Tables S1 to S7

**Other Supplementary Material for this manuscript include (separate file):**

Data S1 [V(D)J sequences of recombinant monoclonal antibodies (Fig. 6A)]

### Methods

#### Patient samples

Peripheral blood mononuclear cells (PBMCs) were isolated from peripheral blood from 58 patients with pediatric peanut allergy (PA) and 13 non-allergic children. Peanut specific IgE (PsIgE) levels obtained on the day of blood draw were used to select the patients. Among 58 pediatric PA, 18 patients had peanut specific IgE higher than 100 kU/L, (PsIgE>100), 13 patients had sIgE lower than 5 kU/L (PsIgE<5) and 24 patients had sIgE between 8.0 to 76.6 kU/L (5<PsIgE<100). Non-allergic children (non-PA) were healthy children without any known history of malignancy, inflammatory or allergic disease, or any known infection at the time of blood collection. The age range and median were similar among cohorts: 5-14 years with median age of 10 for PA children with PsIgE>100; 4-15 years and median age of 9 for PA children with PsIgE<5; 6-17 years and median age of 12 for PA children with 5<PsIgE<100; 4-19 years and median age of 11 for non-allergic children. In terms of incidence of other atopic diseases, 89% of PA children with PsIgE>100 and 69% of PA children with PsIgE<5 reported a history of AD or asthma, which was not significantly different between the two groups (*p=0.147 by chi-square test*, **Table S1**). All patients were screened by board-certified pediatricians and were consented following ISMMS IRB approved protocol.

Peripheral blood was collected in heparin tubes (BD, #367874) and processed within 24 hours. Blood was spun at 1,500 rpm for 10 min. The plasma layer was collected and stored at -80°C. PBMCs were separated by Ficoll-Paque Plus density gradient centrifugation (Sigma-Aldrich, #GE17-1440-02) using Leucosep tube (Grainer Bio-One, #227288) following the manufacturer's instructions. After the isolation, the PBMCs were counted and resuspended in 500 ul of 5% AB serum in RPMI and 500 ul of 20% DMSO in AB serum at 0.5 – 10 x $10^6$ PBMCs per cryovial and cryopreserved.

**Flow cytometry**

Cryopreserved PBMCs were thawed at 37°C and 1 ml of IMDM supplemented with 10% heat inactivated FBS, L-glutamine, sodium pyruvate, non-essential amino acids, and penicillin-streptomycin (complete IMDM) was added dropwise to recover PBMCs. Further, 9 ml of complete IMDM was added gently to resuspend the cells. The cells were rested at 37°C for 15 min, then spun at 1500 rpm for 10 min. Cells were resuspended in 10 ml of pre-warmed HBSS and spun down at 1500 rpm for 10 min. Cells were then resuspended in 10 ml of HBSS and the live cell numbers were determined using trypan blue (Gibco, #15250061). Cells were resuspended in staining buffer (HBSS supplemented with 0.5 % BSA and 1 mM EDTA) at 10 to 20 x $10^6$ cells/ml and stained at 4°C with the following antibodies: Human TruStain FcX^tm (human Fc receptor blocking solution, BioLegend, #422302, lot B313420), Alexa Fluor 647 anti-human CD19 (BioLegend, #302220, lot B260236), FITC anti-human CD27 (BioLegend, #356494, lot B305082), PerCP/Cy5.5 anti-human IgM (BioLegend, #314512, lot B268270), PerCP/Cy5.5 anti-human IgD (BioLegend, #348208, lot B273237), PE/Cyanine7 anti-human IgG Fc (BioLegend, #410722, lot B320675), Brilliant Violet 421 anti-human CD23 (BioLegend, #338522, lot B281791), PE anti-human CD124 (IL-4Ra, BioLegend, #302220, lot B260236), Brilliant Violet 421 anti-human CD19 (BioLegend, #302234, lot B275425), PE/Cyanin7 anti-human IgG (BD Pharmingen, #561298, lot 0073639), PE anti-human IgA (Miltenyi Biotec, #130-113-476, lot 5200306900), APC anti-human IgE (BioLegend, #325508, lot B248126). PerCP/Cy5.5 anti-human IgM (BioLegend, #314512, lot B268270), PE/Cyanin7 anti-human IgG (BD Pharmingen,

#561298, lot 0073639), PE anti-human IgA (Miltenyi Biotec, #130-113-476, lot 5200306900), APC anti-human IgE (BioLegend, #325508, lot B248126) were also used for intracellular staining. Cyto-Fast™ Fix/Perm buffer (BioLegend, #426803) was used for the intracellular staining by following the manufacturer's instruction. All flow cytometric data were acquired on LSRFortessa (BD Biosciences) with automatic compensation and were analyzed using the FlowJo program (Tree Star, Inc.).

**10X Genomics scRNAseq of memory B cells**

B cells were isolated from 17 million to 100 million PBMC from 5 peanut allergic children with sIgE>100 and 3 non-allergic children (**Table S2**) by EasySep™ Human B cell Isolation Kit (STEMCELL technologies, #17954). Hash tagged antibodies were added to 3 million B cells at 1:50 from each sample. B cells from 2 PA subjects were combined in experiment 1. In experiment 2, B cells from 3 PA subjects were combined, and B cells from 3 Non-PA subjects were combined. The cells were then stained with BV421 anti-human CD19 and FITC anti-human CD27 antibodies and $CD19^{+}CD27^{+}$ memory B cell populations were sorted on a FACSAria (BD). After confirming cell number and viability, B cells were run on a Chromium 10X controller using 5' chemistry (10X Genomics) in one lane (experiment 1) or two lanes (experiment 2) with an expected recovery rate of 10,000 cells per lane, according to the manufacturer's instructions. Libraries were generated and run on a HiSeq2500 apparatus.

**10X Genomics scRNAseq processing and analysis**

10X Genomics 5' single cell RNA sequencing data was processed with Cell Ranger version 3.1 and aligned to refdata-cellranger-GRCh38-3.0.0. Gene expression information from all subjects was processed using Seurat v4.1.1(*50*) in R v4.1.0(*50*). To remove apoptotic or lysing cells, cells with ≥10% of RNA transcripts from mitochondrial genes were excluded. To exclude poor quality cells, cells with reads from ≤ 400 features were also removed. To distinguish subjects within each sequencing run, hashtag oligonucleotide information was normalized using the centered log-ratio transformation. The subject of origin for each cell was determined using the default parameters of Seurat's HTODemux function. Only cells associated with a single subject were retained. Read counts were log-normalized using a scaling factor of $10^4$. To account for variability in gene expression, log-normalized read counts were then scaled and centered for each feature. The top 2000 variable genes were then identified using Seurat's "vst" method. V, D, and J genes from the IGH, IGL, and IGK loci were removed so that the properties of the BCR expressed by the cell would remain independent of the cluster it is assigned to. Genes coding for the constant regions of antibody isotypes were also removed except for *IGHE*, which we intended to use as a marker gene. We then used Seurat's IntegrateData function to combine data from the three sequencing runs. Integration was performed using the previously identified top variable genes of each run, and the first 20 dimensions. Following integration, variable gene expression values were re-scaled and centered. This data was then reduced to the first 20 principal components.

B cell subtype identification utilized two clustering steps: initial clustering to identify and remove non-memory B cell contaminants, followed by secondary clustering to resolve memory B cell subtypes. For initial clustering, cells were clustered by Seurat's shared nearest neighbor clustering algorithm with a resolution of 0.25. The B cell subtype of each cluster was then determined by gene expression correlations to cell types in the immunoStates database(*51*). This resulted in 5 clusters identified as memory B cells, and 2 clusters identified as T cells and naive B cells which were removed. Within the remaining memory B cell clusters, data were re-scaled, reduced to the first 100 principal components, and clustered with a resolution of 0.5. This resulted in 10 memory B cell clusters, which were further confirmed to be memory B cells based on expression of marker genes (CD24, TNFRSF13B, **Fig. S6**) and association with mutated BCRs (**Fig. 2D**).

To identify differentially expressed genes in each cluster, the FindAllMarkers function from Seurat was used. Non-integrated gene expression values in each cluster were compared to all other clusters combined. P values were calculated using a Wilcoxon rank-sum test and were adjusted using a Bonferroni adjustment. Genes were classified as differentially expressed within a cluster if their adjusted p values were < 0.05. To identify enrichment of gene ontologies for each cluster, the list of positively differentially expressed genes for each cluster was tested using *enrichR* v3.0(*52, 53*). Differentially expressed genes were compared against the GO Biological Process 2021 database(*52, 53*) within the *enrichR* framework(*54*).

**Memory B cell cultures**

For memory B cell cultures, two distinct $CD19^{+}CD27^{+}$ memory B cell populations were sorted from PBMC: switched $IgM^{-}IgD^{-}IgG^{+}CD23^{-}IL4R^{-}$ (IgG $CD23^{-}IL4R^{-}$ memory), switched $IgM^{-}IgD^{-}IgG^{+}CD23^{+}/IL4R^{+}$ (IgG $CD23^{+}$ memory). To obtain a few thousand B cells per subject for cultures all $CD23^{+}$ and/or $IL4R^{+}$ memory B cells were combined as $CD23^{+}$ cells, since $IL4R^{+}$ memory B cells are at very low frequency, and this receptor is part of the IL-4/IL-13 pathway. Memory B cell populations were sorted in a FACSAria III (BD Biosciences) and seeded in complete IMDM medium at 1000 B cells/well in 96-well plates coated with mitomycin C treated fibroblasts expressing human CD40 and human BAFF (h40LB cells). The cytokines human IL-4 (20 ng/ml, BioLegend, #574002), IL-2 (4 ng/ml, Peprotech, #200-02), IL-10 (10 ng/ml, BioLegend, #571004) and IL-21 (100 ng/ml, BioLegend, #571206) were added at the initiation and on day 4 of culture.

On day 7 of the culture, the supernatants were harvested for ELISA, and B cells were harvested for cell number quantification using counting beads (CountBright Absolute Counting Beads, ThermoFisher Scientific, #C36950) and intracellular staining to evaluate B cell class switching by flow cytometry.

**ELISAs**

To measure total human IgE, Human IgE ELISA$^{BASIC}$ kit (HRP) (MABTECH, #3810) was used following manufacturer's instruction. For peanut specific IgM and IgG ELISAs, plates were coated with crude peanut extract was 5 ug/ml in 0.05M sodium carbonate/bicarbonate buffer, pH 9.6 and incubated overnight at 4°C. Plates were washed 3 times with 0.05% Tween-20 PBS (PBST), blocked with 1 % BSA/PBST at room temperature for 1 hour and washed 3 times with PBST. Undiluted culture supernatants were added to each well and incubated at room temperature for 4 hours or overnight at 4°C. After washing 3 times, biotinylated anti-human IgM at 1:20000 (Jackson ImmunoResearch, #109-066-129) or biotinylated anti-human IgG at 1:40000 (Jackson ImmunoResearch, #109-066-088) were added and plates were incubated at room temperature for 1 hour. The plates were then washed 5 times after which Streptavidin HRP (BD Biosciences, #554066) at 1:1000 was added to the wells. After 30 minutes incubation at room temperature, plates were washed and TMB substrate set (BioLegend, #421101) was added. The reaction was stopped by addition of 0.5 M sulfuric acid. O.D.450 was read in a Molecular Devices SpectraMax apparatus and analyzed by SoftMax Pro (Molecular Devices). For peanut components Ara h 2, Ara h 1 and Ara h 3 ELISAs, purified natural protein (Indoor Biotechnologies) at 2 ug/ml in PBS was used for plate coating.

**PacBio single cell BCR sequencing of Ara h 2 binding cells**

Ara h 2 multimers were generated by associating biotinylated Ara h 2 with PE-labeled streptavidin or AF647-labeled streptavidin. Similarly, detoxified diphtheria toxin (DT) (CRM197, PFENEX Biopharmaceuticals) was used to generate DT multimers. Pre-enriched B cells by EasySep$^{TM}$ Human B cell Isolation Kit (STEMCELL technologies, #17954) from 8 PA with PsIgE>100, 5 PA with PsIgE<5 and 2 non-allergic subjects were used for Ara h 2 binder and/or DT binder sorting (Table S5). Single Ara h 2 (or DT) binding B cell was sorted using FACSAria III (BD Biosciences) into each well in the 96 well plates with lysis buffer containing RNAse inhibitor, dNTPs and oligo dT-VN30 anchored with template switch oligo sequence (**Table S6**). Plates were flash frozen in dry ice and stored at -80°C. Plates were thawed on ice, then oligo dT was added and annealing was performed at 70°C for 5 min. cDNA synthesis and cDNA amplification was performed with template switching oligo (TSO) using template switching RT enzyme mix (NEB) according to the manufacturer's instruction. The first PCR was performed to amplify BCR heavy and light chains using outer primers targeting human immunoglobulin constant regions (IgG, IgM for heavy chains, kappa and lambda for light chains) and TSO inner primer (**Table S6**), respectively. After cleaning up the PCR products, BCR heavy and light chains were further amplified with inner primers with plate and row specific ID (5') and column specific ID (3') to enable the identification of BCR sequences derived from each well (**Table S6**). Afterwards, all PCR products were combined as one library. PacBio sequencing was performed to obtain long amplicon reads to cover 500 to 1000 bp. All PCR reactions were performed using NEBNext Ultra$^{TM}$ II Q5 master mix (NEB).

**Analysis of gene expression using real-time PCR**

Quantification of *CD23*, *GLT-IGHE*, and *total-IGHE* expression in IgG1 clones from antigen-binding cells were performed in an Applied Biosystems™ QuantStudio™ 6 Pro Real-Time PCR System (Thermo Fisher Scientific) with LUNA™ Universal qPCR Master Mix (NEB), according to the manufacturer's instructions. The cDNAs from single B cells derived from Ara h 2 binders (n=34) from 6 PA subjects with PsIgE>100 and DT binders from 6 PA subjects with PsIgE>100 (n=14) and 2 non-allergic children (Non-PA, n=13) were analyzed. The mRNA relative abundances were normalized to β-actin (as the reference gene) using the cycle threshold (Ct) method. The sequences of all reported and self-designed primers are listed in **Table S7**.

**B cell receptor sequence processing and analysis**

BCR sequencing was obtained from two sources: 10X Genomics scRNA-seq + BCR, as well as sorted single cell PacBio sequencing. BCRs from both were processed using the Immcantation suite (immcantation.org). BCR sequence data from 10X Genomics scRNA-seq + BCR sequencing began with the filtered V(D)J contigs from 10X Genomics Cell Ranger version 3.1. To obtain V and J gene assignments, these contigs were aligned to the IMGT GENE-DB germline reference allele database (obtained 8/3/2019)(*55*). Non-productive heavy and light chain BCR sequences were removed. BCRs were annotated by B cell subtype based on matching their single cell barcodes to the gene expression information. Only cells that had both gene expression and BCR sequence data were retained. In cells with multiple heavy chains, only one heavy chain with the highest unique molecular identifier count was retained. In the event of a tie, the first heavy chain identified was retained.

For BCR sequencing data from sorted single cell PacBio sequencing data, adaptor ligation, sequencing reactions, and initial sequence processing were conducted at GENEWIZ, LLC/Azenta

US, Inc (South Plainfield, NJ, USA) using standard PacBio circular consensus sequence (CCS) tools and guidelines. Further processing was performed using *presto* v0.6.2(*56*). To remove reads too short to likely contain functional BCRs and associated constant regions, reads shorter than 500bp were removed. To identify the plate and row of each read, forward primers were aligned to the first 1000bp of each read, with upstream sequence sites removed. To identify *IGHG* sub-isotype sequences, internal constant region sequences associated with each subisotype were aligned to each read. Reads were tagged if they contained a match to an *IGHG* sub-isotype constant region with a maximum error rate of 0.1. To identify plate column and constant region primer (IgG, IgM, IgK, IgL) for each read, reverse primer sequences were also aligned to each read. Primer sequences are available in **Table S6**. Nucleotide sites matching primer sequences, as well as the sites downstream of them, were removed. Reads failing to exactly match both a forward and reverse primer were discarded. Remaining reads shorter than 500bp in length were also discarded. After initial processing, PacBio BCR sequence reads were assembled into consensus sequences. Because each well within each plate should represent reads from a single B cell, reads within a forward and reverse primer combination should correspond to a single consensus sequence. However, initial analyses indicated further filtering was required to obtain consensus sequences with appropriate confidence. Within each forward and reverse primer combination, reads were further clustered using *vsearch* v2.14.1(*57*) with an 80% similarity threshold. To account for indel variation within a cluster, reads within these clusters were aligned to each other using *MUSCLE* v3.8.1551(*58*). Reads within each of these clusters were assembled into consensus sequences. Nucleotide sites in which < 90% of reads had the same character, or had a consensus quality score < 20, were assigned as ambiguous "N" characters. Sites in which >50% of reads contained gap characters were removed. Read groups containing less than 10 reads or in which >10% of sites differed from their final consensus sequence were removed. As with 10X Genomics BCR sequence data, consensus sequences were aligned to the IMGT GENE-DB database(*55*) to obtain V and J germline annotations. Non-productive heavy and light chain BCR sequences were removed. 18.3% of forward/reverse primer combinations yielded multiple productive consensus sequences from different clusters. These sequences possibly represent doublets and were discarded unless one sequence, which was retained, accounted for at least 90% of the reads in that well/constant region group. For each consensus sequence, *IGHG* subisotype was determined as the most frequent-matching subisotype sequence among constituent reads, or "Unknown" if no subisotype sequence match was found. Confirming the accuracy of this step, *IGHG* subisotypes could only be determined for sequences that also matched *IGHG* primers. As further quality control, 16 sequences originated from wells in which no Ig DNA was detected through PCR and were discarded. Additionally, 13 sequences with < 300 non-ambiguous nucleotides were also discarded. This pipeline yielded 243 total heavy chain sequences, 210 of which were paired with functional light chain BCRs. We validated our consensus sequences through two tests. First, consensus sequences assembled with high error rates are unlikely to exactly match IMGT reference sequences. We tested this in our data by quantifying the SHM frequency of *IGHM* heavy chain consensus sequences. While sequences from *IGHG* primers were highly mutated (median SHM = 8.7%), most *IGHM* consensus sequences were mostly unmutated (median SHM = 0%, **Fig. 5A**). Next, heavy and light chain sequences assembled from different B cells are unlikely to have similar levels of SHM. In our consensus sequences, however, SHM levels of paired heavy and light chains were closely correlated (linear regression slope $p<2e\text{-}16$, $R^2=0.63$).

For BCR data from both data sources, SHM level was determined for each cell as the frequency of non-ambiguous mismatches along the IGHV gene of the unmutated germline sequence along

IMGT positions 1-312 using *shazam* v1.1.2(*59*). To identify potential Ara h 2-binding convergent sequence families across all single cell datasets, sequences from all datasets were partitioned based on common IGHV and IGHJ gene annotations, as well as junction region length. Within these groups, sequences differing from one another by an amino acid Hamming distance threshold of 0.2 within the junction region were clustered together using single linkage hierarchical clustering(*60*) implemented in *scoper* v1.2.0(*61*). When searching against public sequences, we also identified one cell (P4AC5) with a matching V gene, J gene, and junction amino acid Hamming distance of 0.21 compared to P413P1E10. This cell was also included within the P413P1E10 family and confirmed to bind to Ara h 2 through ELISA (**Fig. 6B**). In the 4C5G1 convergent sequence family, all cells expressed similar IGK light chains, however cell P1DC5 also expressed an IGL light chain (IGLV4-69*01/IGLJ3*02). This sequence was not included in further analyses. Consensus unmutated V and J gene segments for each convergent sequence family (**Fig. 6A**) were constructed using the createGermlines function in *dowser* v1.1.0 (*62*). For ambiguous V gene assignments, only the first assignment was used for reconstruction.

**Recombinant antibody expression**

Selected BCR heavy and light chain V(D)J gene sequences were used to generate recombinant antibodies to verify binding specificity and affinity. All heavy chains were expressed as human IgG1, and light chains were expressed either as lambda or kappa according to the original BCR usage. Gene synthesis and recombinant antibody production were performed by Twist Biosciences. All antibody sequences tested are available in the file Data S1.tsv.

**Biolayer interferometry**

To evaluate kinetic interactions of the recombinant antibodies with peanut proteins, biolayer interferometry was performed on a ForteBio Octet 96 following the manufacturer's instruction. Anti-human IgG Fc capture biosensors (AHC) were used to capture recombinant antibodies with 1x PBS as the assay buffer. Purified natural Ara h 1, Ara h 2 and Ara h 3 (were used at 117.6, 58.8, 29.4, 14.7, 7.35, 3.68 and 1.84 nM to measure the antibody-antigen interaction. Purified antibodies were loaded at a concentration of 10 ug/ml. The following protocol was used for each assay: 60 s baseline, 300 s antibody load, 120 s baseline2, 300 s association with protein, 900 s dissociation. Before the start of each run, fresh biosensors were activated via soaking in PBS for 10 minutes. Data from each assay was processed using ForteBio software. Curves were analyzed using a 1:1 binding model, fit globally between time 115 seconds and 120 seconds after subtraction of the ligand reference sensors.

**Statistical analysis**

Unless otherwise specified, data were analyzed and graphed with GraphPad Prism 9 software (GraphPad Software). The *Man-Whitney U test* was used to compare medians between groups. The *Spearman ranked correlation coefficient* was used to assess the significance of correlations between IgE levels and specific IgE levels. *Chi-square test* was used to determine the significant differences between expected frequencies and observed frequencies.

**References** 50-62

### Supplementary Figures and Supplementary Tables

**Fig. S1**

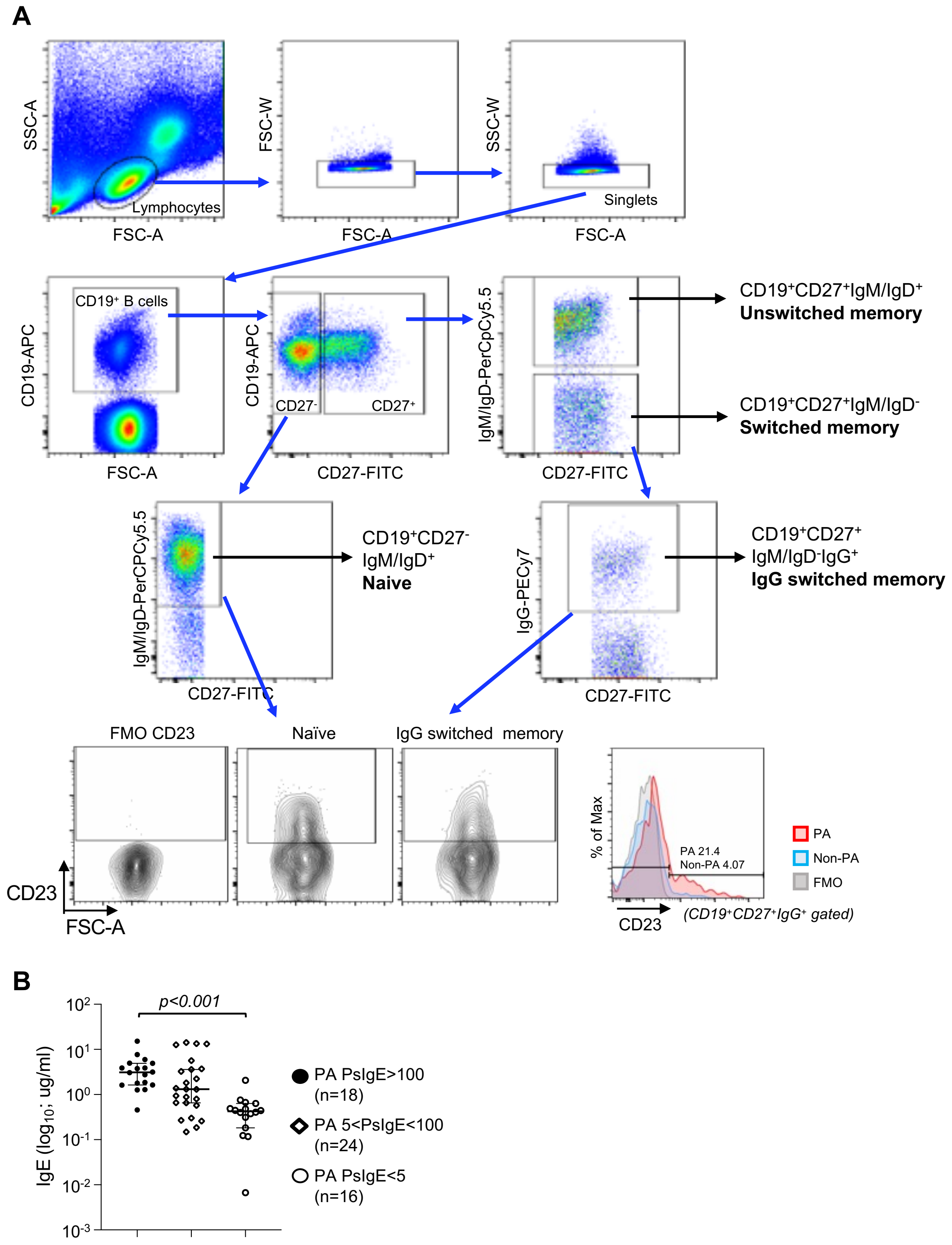


**Fig. S1. Gating strategy to analyze memory B cell populations in PBMC by flow cytometry.**

**(A)** Gating strategy to identify memory B cell population for flow cytometry. **(B)** Total plasma

IgE levels in PA PsIgE>100 was higher than those in PA with PsIgE<5 (*$p<0.0001$ by Mann Whiteney U test*). Red solid bar; PsIgE>100 PA, white open bar: PA with PsIgE level between 5 to 100, red shaded bar; PsIgE<5 PA.

**Fig. S2**

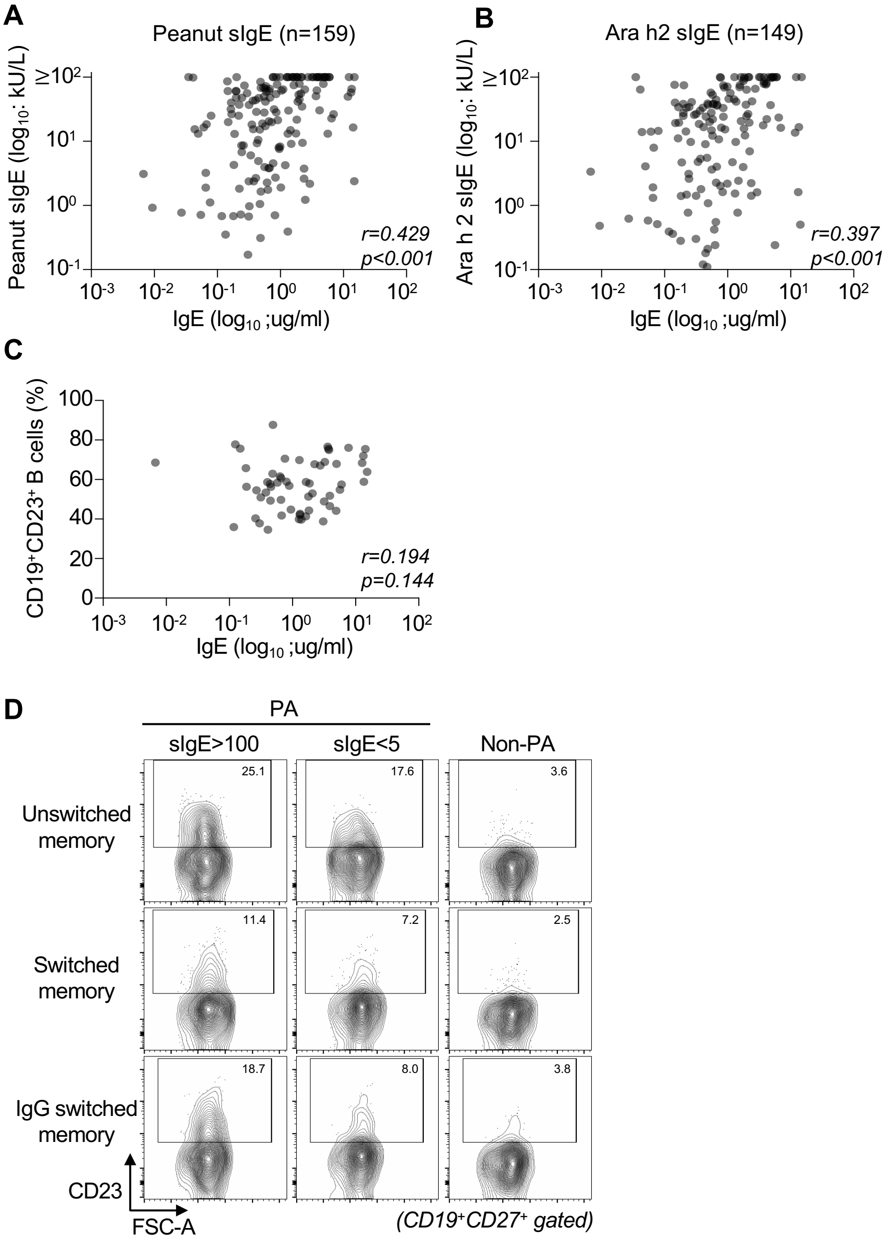


**Fig. S2. Correlation analysis of antigen specific IgE and total IgE level.** (**A**) Peanut specific IgE and (**B**) Ara h 2 specific IgE were positively correlated with total IgE level. (**C**)The frequency of total $CD19^{+}CD23^{+}$ B cells was not correlated with total IgE in PA children (n=58). *Spearman ranked correlation coefficient* was used for statistical analysis. (D) Representative data of CD23 staining among memory B cell populations from PsIgE>100 PA (left), PsIgE<5 PA (center) and Non-PA children (right) are shown.

**Fig. S3**

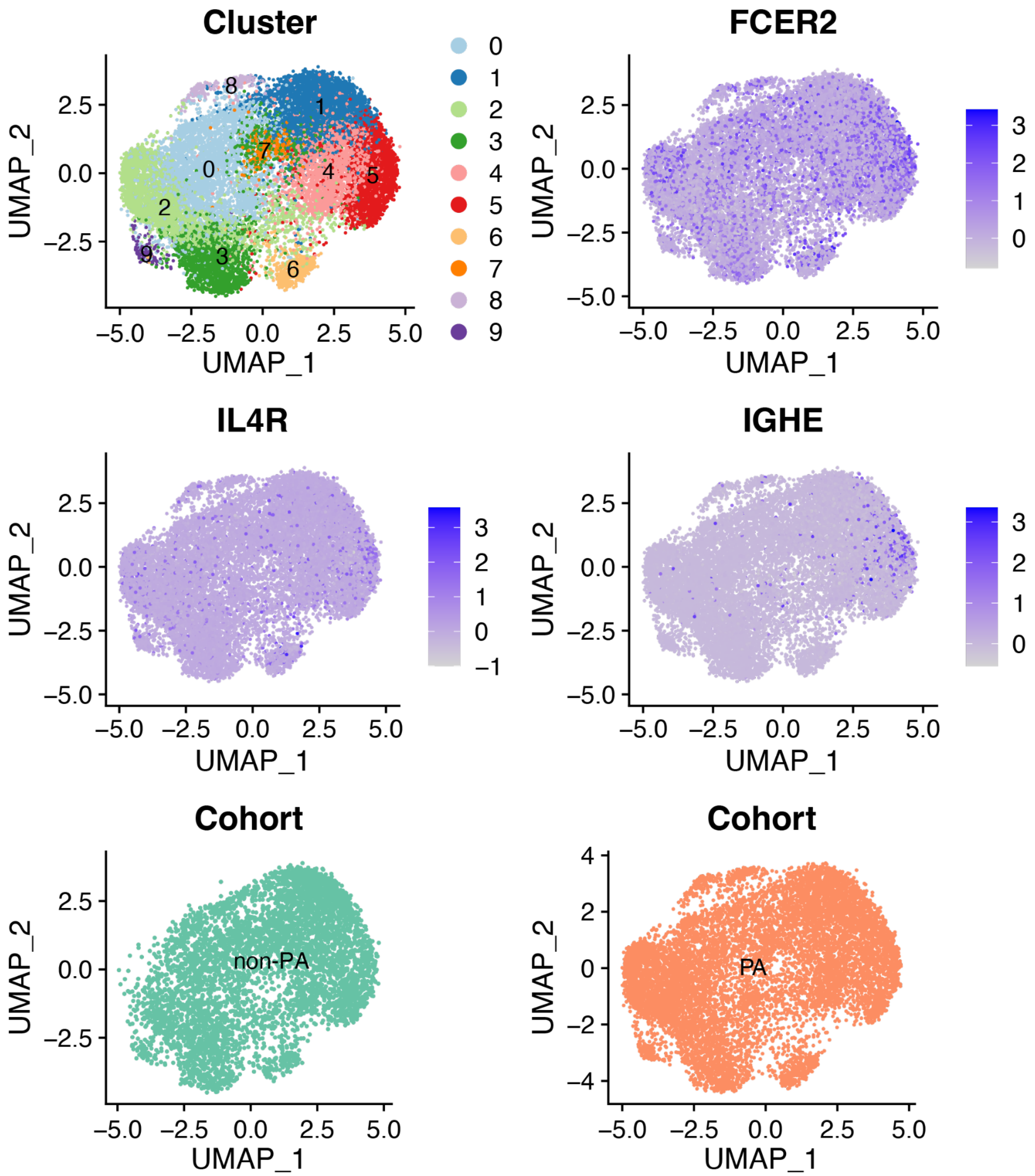


**Fig. S3. UMAP colored by gene cluster, gene expression, and cohort.** Each dot represents a single cell. These cells are colored by normalized expression of *FCER2*, *ILR4*, and *IGHE*. In the bottom two panels, cells are colored by cohort. Cells colored by cluster are shown in the top left (same as **Fig. 2A**) for reference.

**Fig. S4**

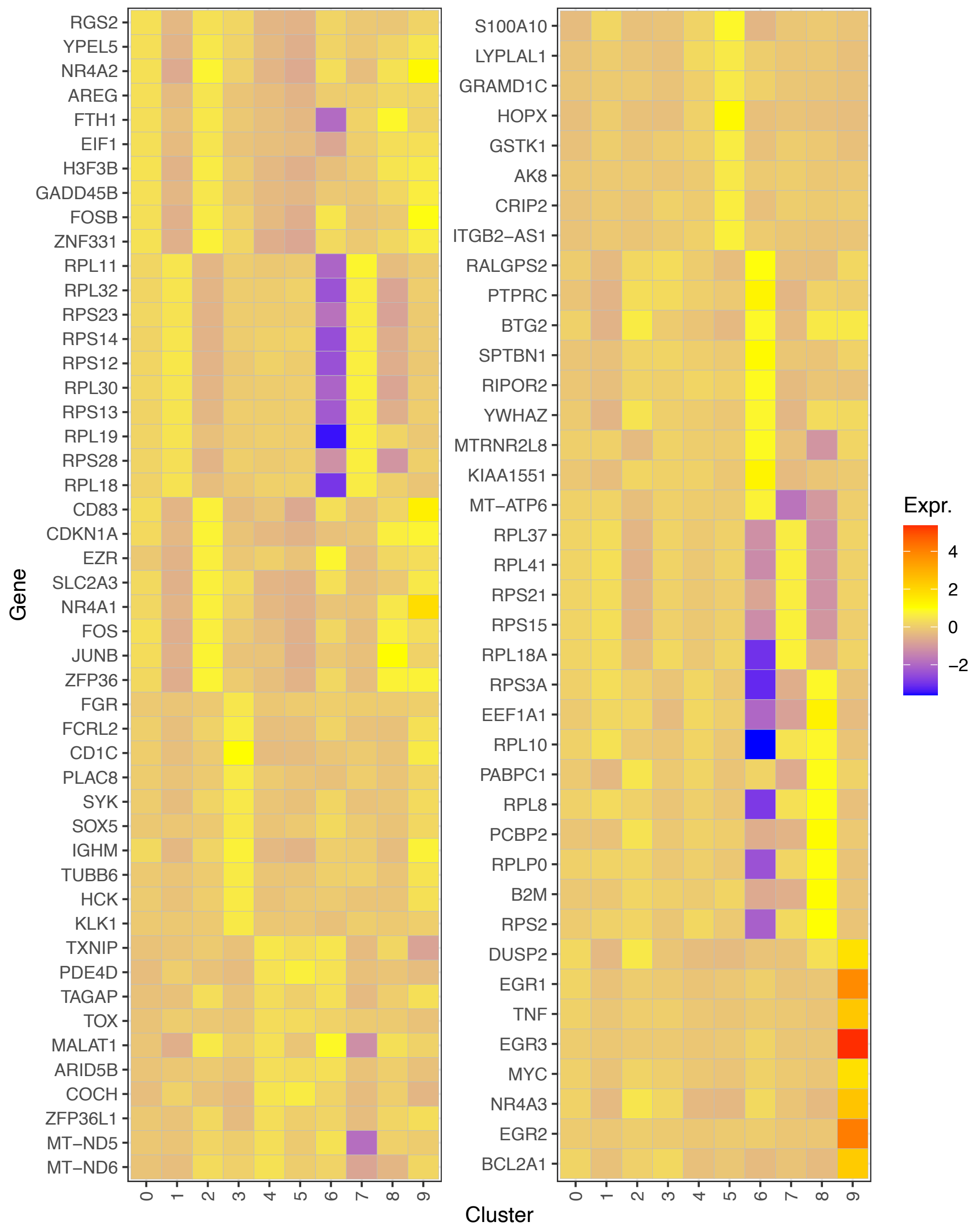


**Fig. S4. Scaled expression of top differentially expressed genes for cells in all clusters.** Similar to **Fig. 2C**, but scaled, normalized expression for all clusters. Only the top 10 positively differentially expressed genes for each cluster, minus duplicates, were included.

Fig. S5

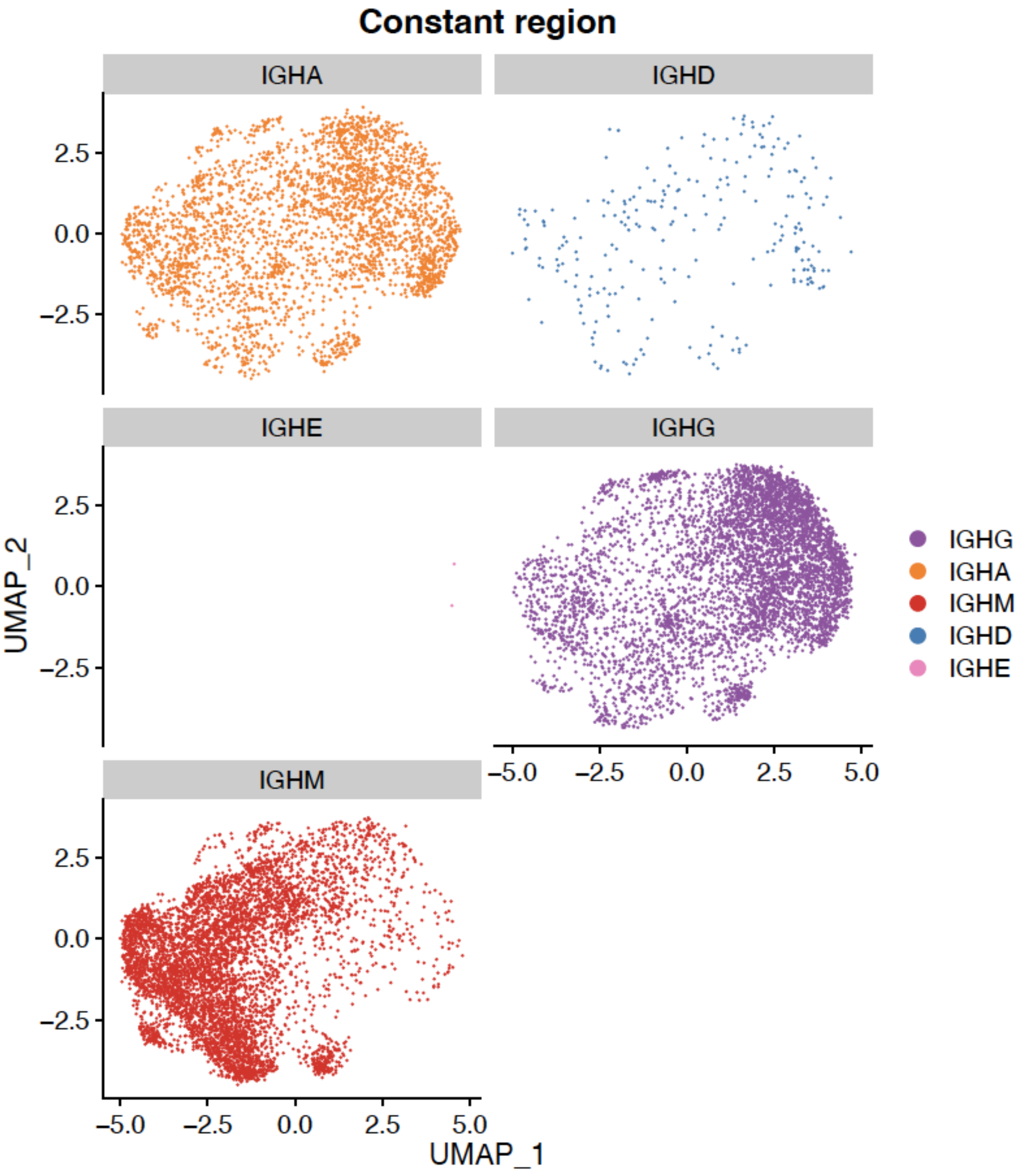


**Fig. S5. UMAP colored by heavy chain constant regions.** Corresponds to the scRNAseq analysis in **Fig. 2** and **Fig. 3**.

**Fig. S6**

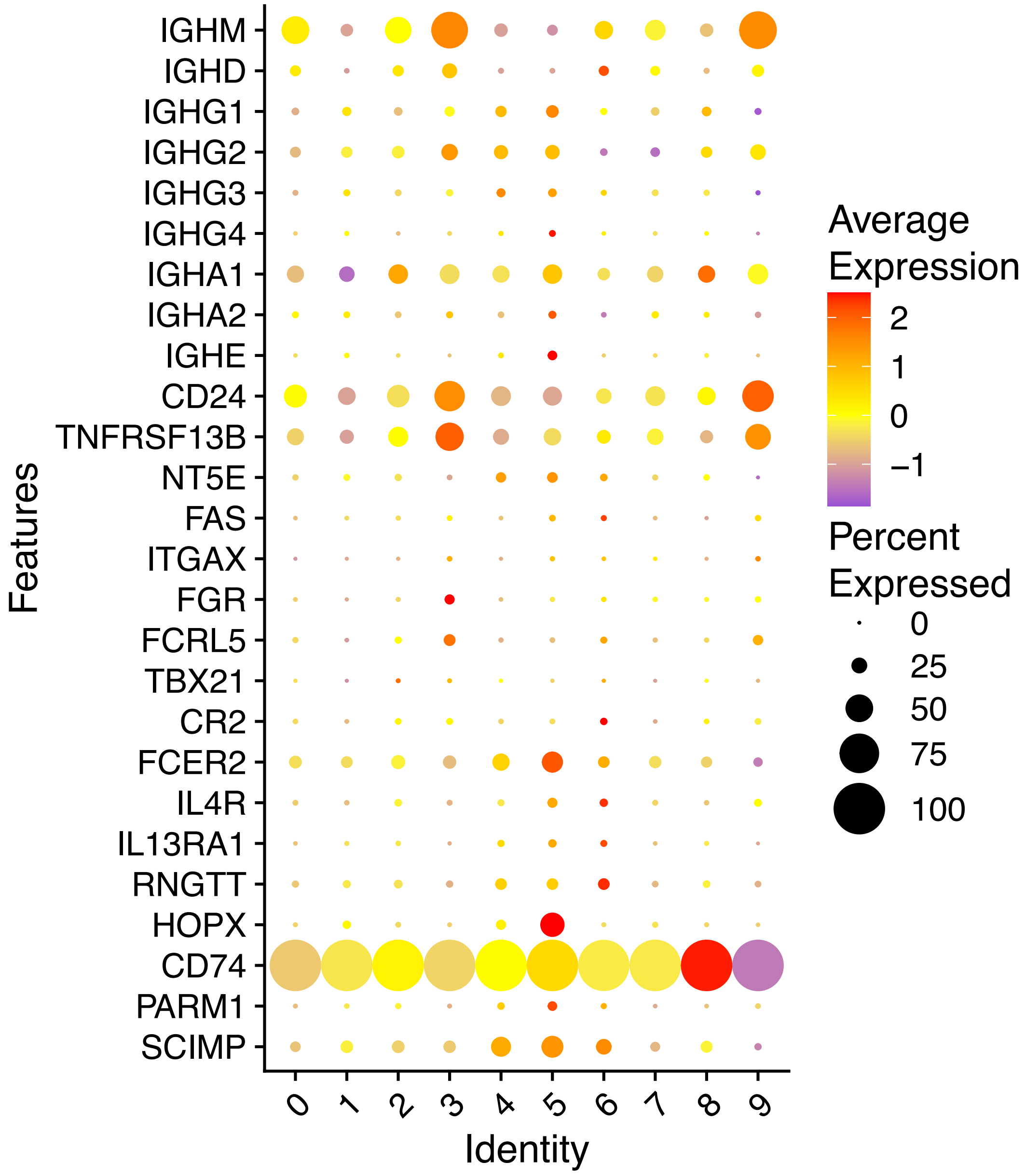


**Fig. S6. DotPlot showing memory B cell marker genes.** Corresponds to the scRNAseq analysis in **Fig. 2**.

Fig. S7

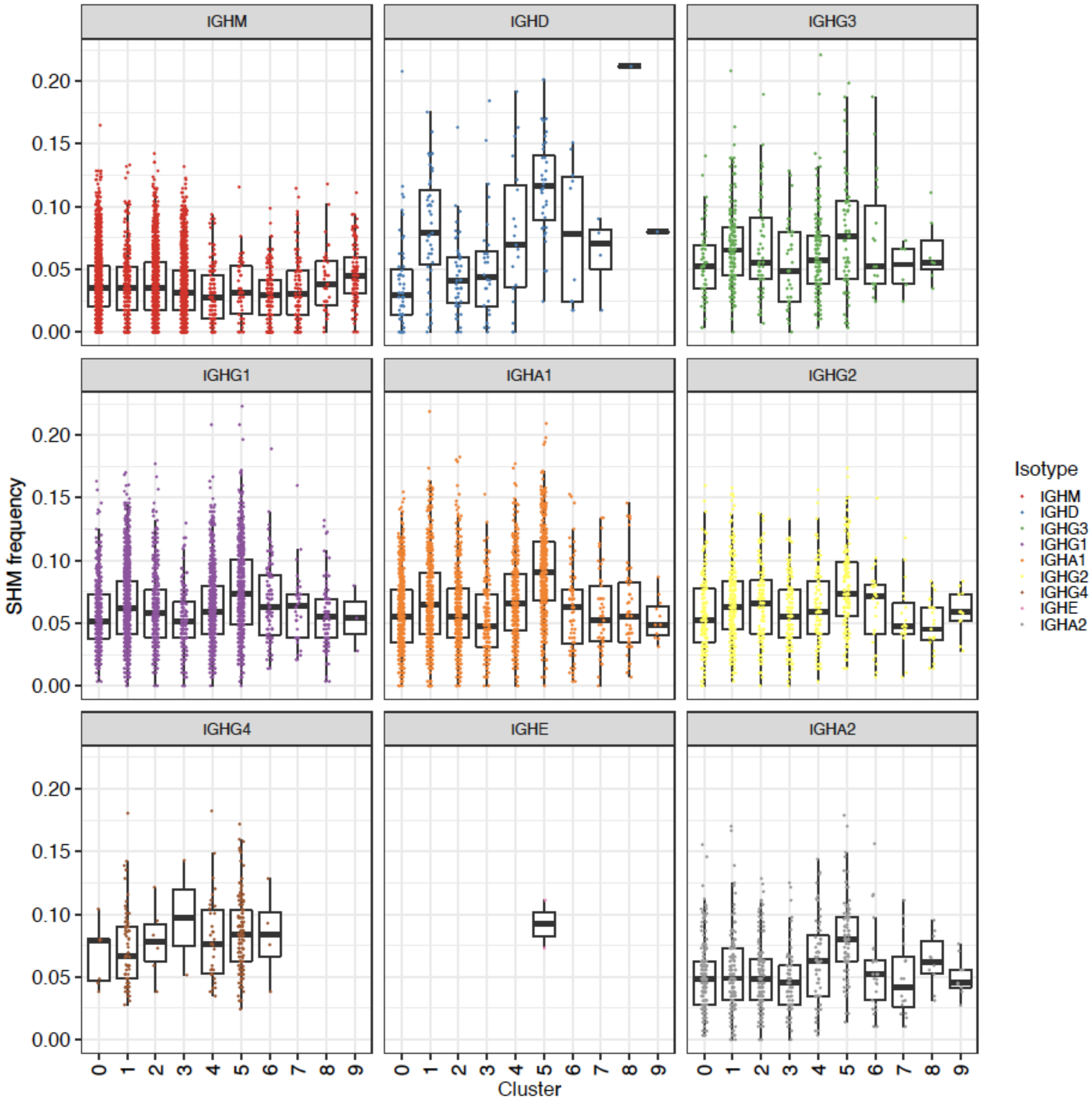


**Fig. S7. Somatic hypermutation levels of B cells in each cluster, colored by BCR constant region.** Corresponds to the scRNAseq analysis in **Fig. 2** and **Fig. 3**.

**Fig. S8**

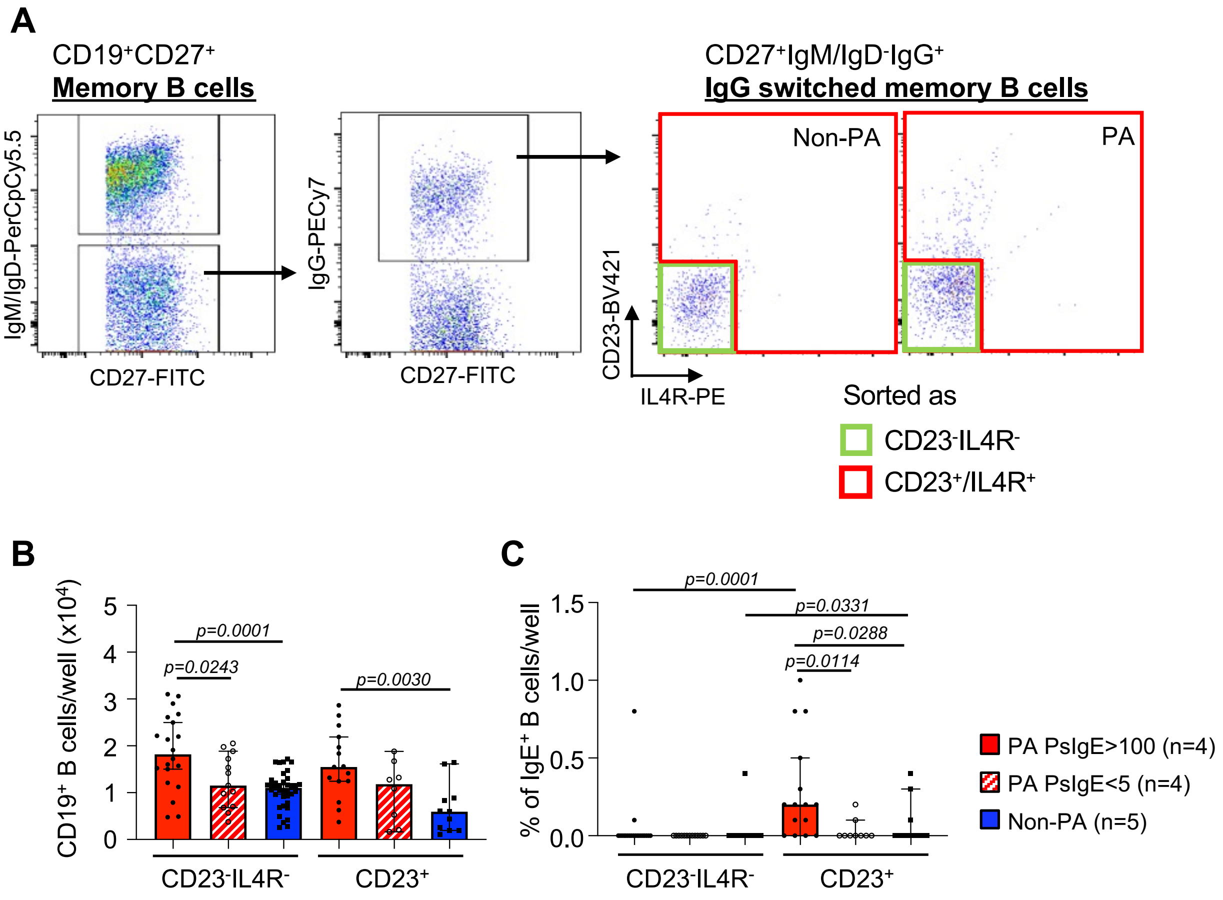


**Fig. S8. CD23$^+$IgG$^+$ switched memory B cell culture from PA children with PsIgE>100 showed more proliferation and more class switching to IgE in vitro. (A)** Gating strategy for sorting IgG switched memory B cells from PBMC. **(B)** CD23$^+$IgG$^+$ switched memory B cells from PA with PsIgE>100 showed more proliferation than non-allergic children (*p=0.0030, by*

*Mann-whiteny U test*) **(C)** More CD23$^{+}$IgG switched memory B cells from PA with PsIgE>100 PsIGE>100 class switched to IgE in vitro on day 7 of the culture even with IL-21. (*p=0.0114* vs PA with PsIgE<5, *p=0.0288* vs non-allergic children by *Mann-whiteny U test*).

**Fig. S9**

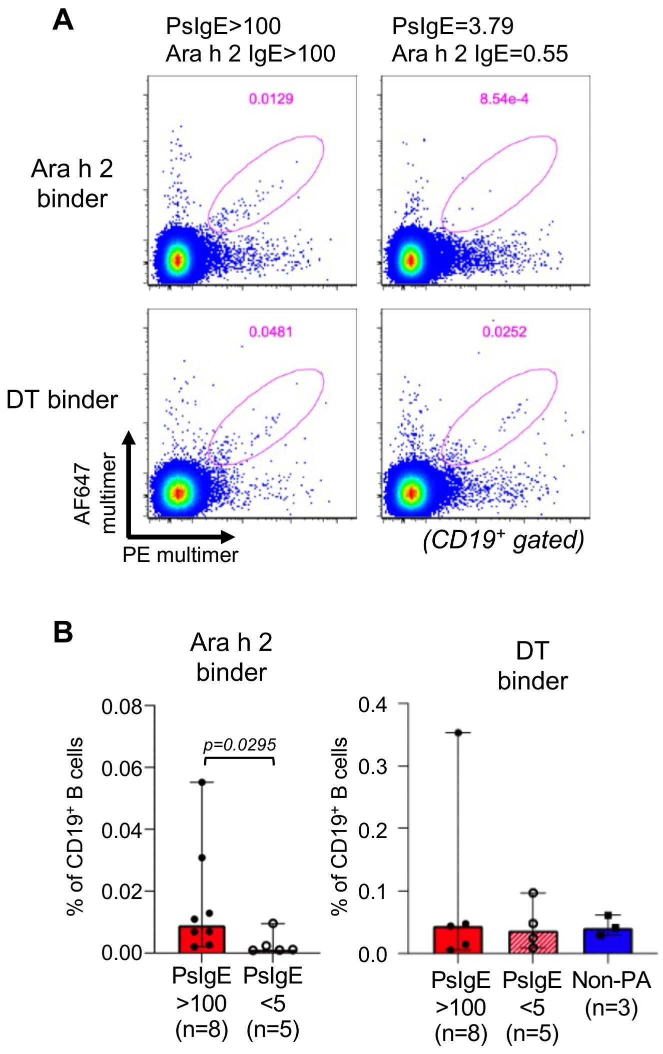


**Fig. S9. Detection of Ara h 2 or diphtheria toxin (DT) binding B cells. (A)** Pre-enriched B cells were stained with AF647 and PE labeled multimers of Ara h 2 (top panel) or DT (bottom panel) to detect antigen binding B cells in PsIgE>100 and PsIgE<5 PA. **(B)** The frequency of Ara h 2 binder cells was higher in PsIgE>100 than PsIgE<5 PA (*p=0.0295, Mann-Whitney U test*) while there was no significant difference in the frequency of DT binders among PsIgE>100, PsIgE<5 and non-PA children **(C)**.

**Fig. S10**

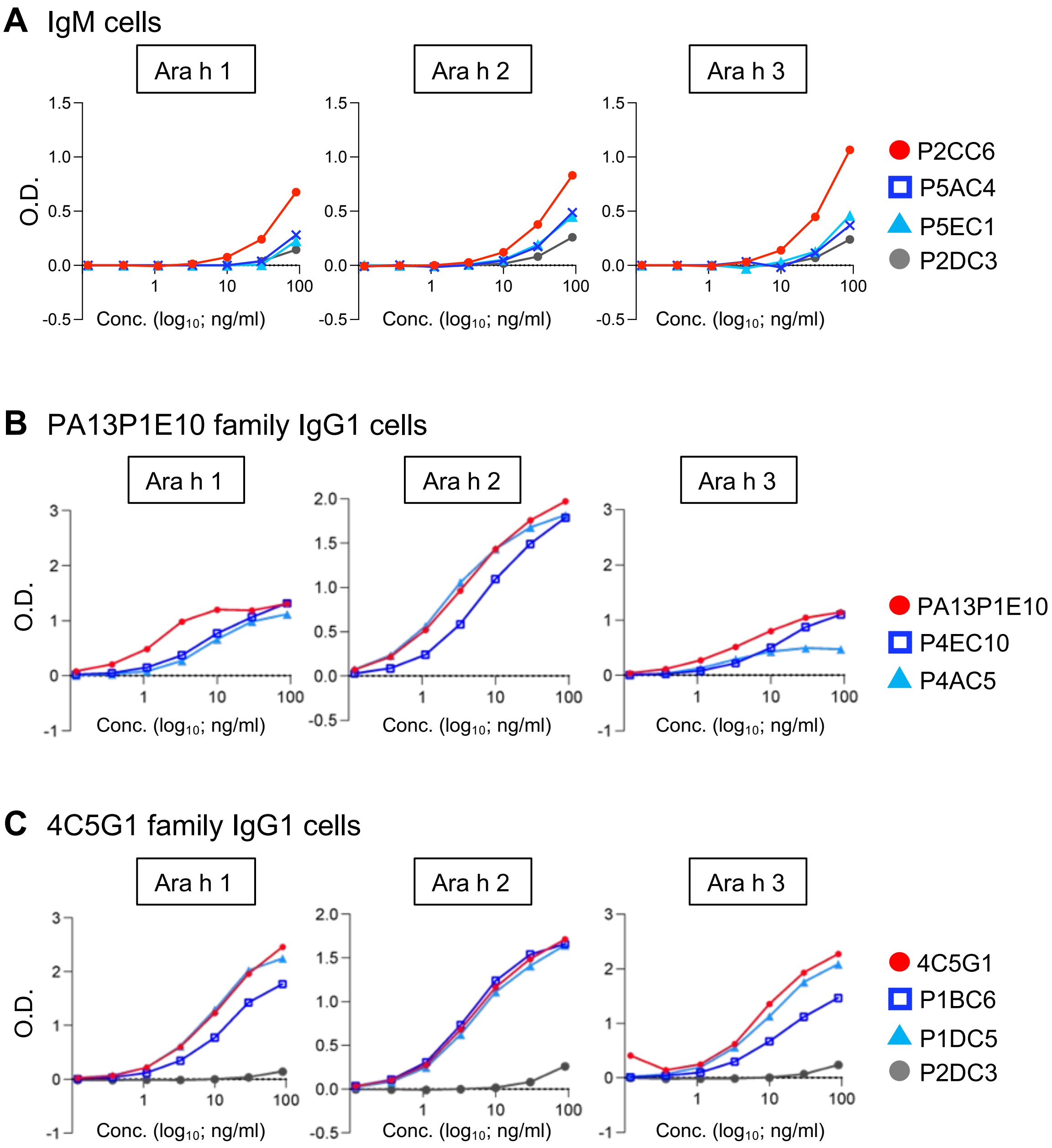


**Fig. S10. Binding properties of recombinant antibodies against Ara h 1, 2 and 3 by ELISA.** Antibodies from **(A)** three IgM cells among Ara h 2 binders were expressed as IgG1 to measure the binding against Ara h 1, 2 and 3 by ELISA. **(B)** IGHV3-30/IGKV3-20, which are highly similar to PA13P1E10 and **(C)** IGHV1-8 convergent sequence families found in this study are tested for the binding by ELISA. DT binding IgG1 cell P2DC3, which used IGHV1-8, was used as a negative control.

**Fig. S11**

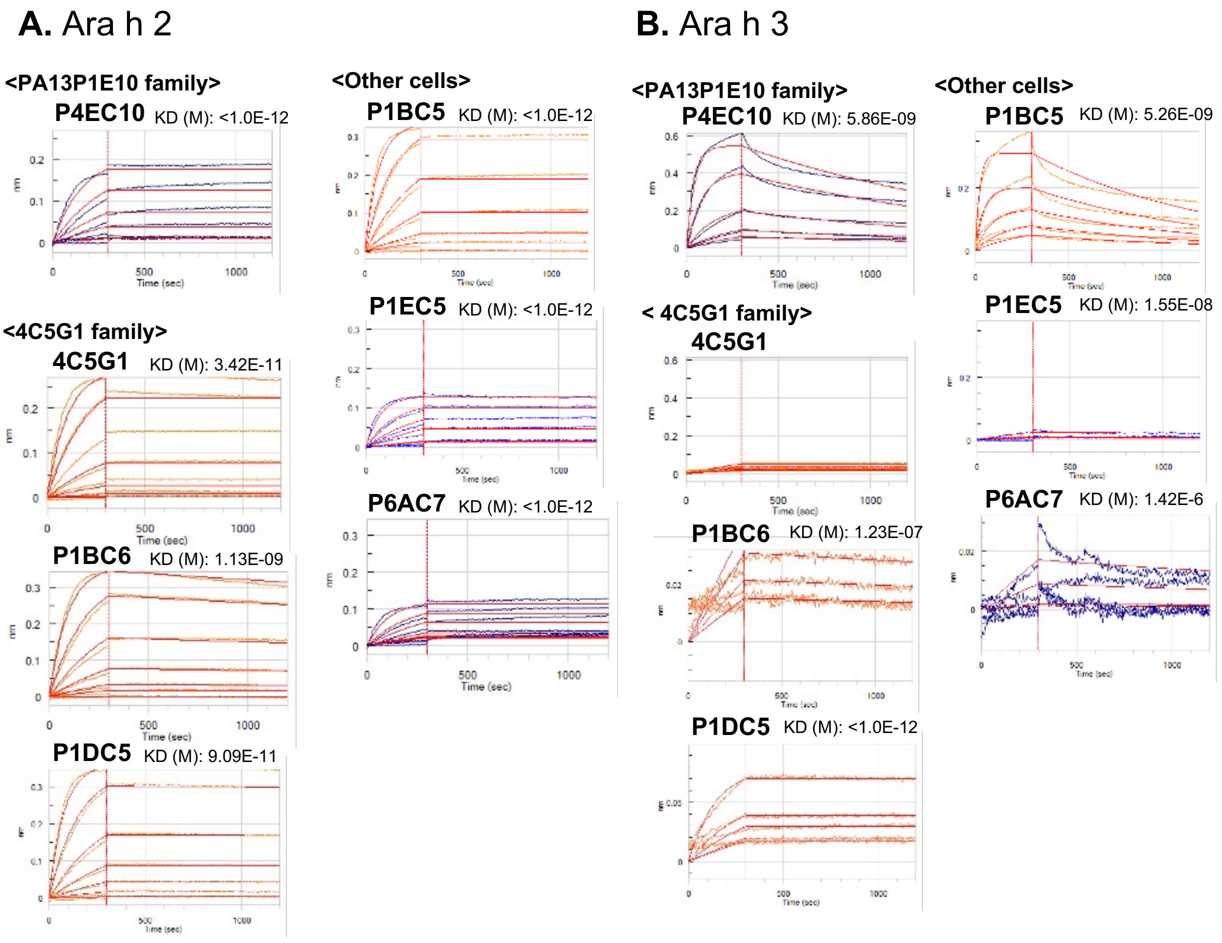


**Fig. S11. Affinity measurement of recombinant anti-Ara h 2 antibodies by biolayer interferometry.** The binding affinity was measured by association and dissociation of antibody and Ara h 2 **(A)** or Ara h 3 **(B)** interactions. Each obtained KD (M) is indicated.

**Table S1.**
Demographics of samples used in flow cytometry analysis.

| Sample ID | Sex | Age | Allergic status | total IgE (ug/ml) | Peanut sIgE (kU/L) | Ara h2 sIgE (kU/L) | AD | Asthma |
|---|---|---|---|---|---|---|---|---|
| **H1** | F | 10 | Allergic | 0.454 | >100 | n.a. | No | No |
| **H2** | F | 5 | Allergic | 4.898 | >100 | n.a. | No | Yes |
| **H3** | M | 11 | Allergic | 1.633 | >100 | n.a. | Yes | Yes |
| **H4** | M | 6 | Allergic | 7.695 | >100 | n.a. | Yes | Yes |
| **H5** | M | 8 | Allergic | 3.888 | >100 | n.a. | Yes | No |
| **H6** | M | 6 | Allergic | 3.055 | >100 | n.a. | Yes | Yes |
| **H7** | F | 8 | Allergic | 3.162 | >100 | >100 | Yes | No |
| **H8** | F | 13 | Allergic | 1.852 | >100 | 95 | No | Yes |
| **H9** | M | 10 | Allergic | 1.620 | >100 | >100 | Yes | No |
| **H10** | F | 12 | Allergic | 3.890 | >100 | >100 | No | No |
| **H11** | F | 14 | Allergic | 1.283 | >100 | >100 | No | Yes |
| **H12** | F | 9 | Allergic | 3.763 | >100 | >100 | No | Yes |
| **H13** | F | 10 | Allergic | 5.964 | >100 | >100 | Yes | Yes |
| **H14** | F | 7 | Allergic | 1.781 | >100 | >100 | Yes | Yes |
| **H15** | F | 12 | Allergic | 2.739 | >100 | 65.8 | Yes | Yes |
| **H16** | M | 10 | Allergic | 4.968 | >100 | >100 | No | Yes |
| **H17** | M | 11 | Allergic | 1.264 | >100 | 45 | Yes | Yes |
| **H18** | F | 11 | Allergic | 15.167 | >100 | >100 | Yes | Yes |
| **L1** | F | 15 | Allergic | 0.442 | 2.11 | n.a. | Yes | Yes |
| **L2** | F | 4 | Allergic | 0.124 | 0.61 | n.a. | Yes | |
| **L3** | M | 9 | Allergic | 0.482 | 3.78 | n.a. | Yes | Yes |
| **L4** | M | 10 | Allergic | 0.400 | 3.13 | n.a. | No | Yes |
| **L5** | M | 8 | Allergic | 0.670 | 3.83 | 4.01 | No | No |
| **L6** | F | 12 | Allergic | 0.488 | 1.64 | 1.08 | No | No |
| **L7** | M | 14 | Allergic | 0.756 | 3.79 | 0.55 | No | No |
| **L8** | M | 8 | Allergic | 0.182 | 4.27 | 1.4 | | Yes |
| **L9** | F | 12 | Allergic | 0.007 | 3.09 | 3.35 | Yes | No |
| **L10** | M | 9 | Allergic | 0.117 | 0.68 | 0.39 | No | Yes |
| **L11** | F | 7 | Allergic | 0.446 | 4.91 | 0.18 | Yes | No |

| | | | | | | | | |
|---|---|---|---|---|---|---|---|---|
| **L12** | F | 9 | Allergic | 2.064 | 1.22 | <0.1 | No | Yes |
| **L13** | F | 15 | Allergic | 0.632 | 1.75 | 0.24 | No | No |
| **L14** | M | 9 | Allergic | 0.406 | 2.67 | 0.33 | Yes | No |
| **L15** | M | 9 | Allergic | 0.313 | 0.67 | 0.3 | Yes | Yes |
| **L16** | M | 10 | Allergic | 0.377 | 2.37 | 1.79 | | |
| **M1** | F | 12 | Allergic | 3.240 | 9.82 | n.a. | Yes | Yes |
| **M2** | M | 9 | Allergic | 12.653 | 18.7 | n.a. | Yes | No |
| **M3** | F | 15 | Allergic | 1.315 | 35.2 | n.a. | No | Yes |
| **M4** | M | 14 | Allergic | 0.656 | 48.9 | n.a. | Yes | Yes |
| **M5** | M | 11 | Allergic | 0.186 | 64.6 | n.a. | Yes | Yes |
| **M6** | M | 17 | Allergic | 0.940 | 8 | <0.1 | Yes | Yes |
| **M7** | M | 13 | Allergic | 2.231 | 9.44 | 6.96 | | Yes |
| **M8** | F | 13 | Allergic | 5.624 | 13.1 | 0.24 | | No |
| **M9** | F | 7 | Allergic | 1.814 | 21.7 | 15.8 | Yes | Yes |
| **M10** | M | 15 | Allergic | 1.307 | 13.1 | 1.53 | Yes | No |
| **M11** | M | 7 | Allergic | 0.654 | 60.6 | 44.4 | Yes | Yes |
| **M12** | M | 14 | Allergic | 0.884 | 76.6 | 68.7 | Yes | No |
| **M13** | M | 17 | Allergic | 0.575 | 65.8 | 52.9 | | |
| **M14** | F | 11 | Allergic | 3.622 | 14.1 | n.a. | No | Yes |
| **M15** | M | 11 | Allergic | 3.694 | 25.8 | n.a. | Yes | Yes |
| **M16** | | 6 | Allergic | 0.758 | 41.7 | n.a. | | Yes |
| **M17** | F | 14 | Allergic | 0.332 | 58.7 | n.a. | No | Yes |
| **M18** | M | 6 | Allergic | 1.377 | 92 | n.a. | | |
| **M19** | M | 10 | Allergic | 14.201 | 16.4 | 0.5 | No | Yes |
| **M20** | M | 7 | Allergic | 13.152 | 57.6 | 1.61 | Yes | Yes |
| **M21** | M | 15 | Allergic | 0.148 | 19.9 | 19.2 | | |
| **M22** | M | 12 | Allergic | 13.458 | 65.5 | 16.6 | Yes | Yes |
| **M23** | F | 11 | Allergic | 0.268 | 73.9 | 40.8 | Yes | Yes |
| **M24** | F | 15 | Allergic | 0.259 | 8.61 | 9.74 | | |
| **NA1** | F | 4 | Non-allergic | n.a. | n.a. | n.a. | No | No |
| **NA2** | F | 12 | Non-allergic | 0.253 | n.a. | n.a. | No | No |

| | | | | | | | | |
|---|---|---|---|---|---|---|---|---|
| **NA3** | F | 11 | Non-allergic | 0.045 | n.a. | n.a. | No | No |
| **NA4** | F | 16 | Non-allergic | 0.019 | n.a. | n.a. | No | No |
| **NA5** | F | 10 | Non-allergic | 0.020 | n.a. | n.a. | No | No |
| **NA6** | F | 19 | Non-allergic | 0.002 | n.a. | n.a. | No | No |
| **NA7** | M | 11 | Non-allergic | 0.037 | n.a. | n.a. | No | No |
| **NA8** | F | 10 | Non-allergic | 0.360 | n.a. | n.a. | No | No |
| **NA9** | F | 14 | Non-allergic | 0.302 | n.a. | n.a. | No | No |
| **NA10** | F | 14 | Non-allergic | 0.167 | n.a. | n.a. | No | No |
| **NA11** | M | 15 | Non-allergic | 0.079 | n.a. | n.a. | No | No |
| **NA12** | M | 9 | Non-allergic | 0.124 | n.a. | n.a. | No | No |
| **NA13** | M | 9 | Non-allergic | 0.133 | n.a. | n.a. | No | No |

**H**: PA children with PsIgE>100, **L**: PA children with PsIgE<5, **NA**: Non-allergic children
n.a.: not available

**Table S2.**

Demographics of samples used in scRNA seq.

| Sample ID | Sex | Age | Allergic status | total IgE (ug/ml) | Peanut sIgE (kU/L) | Ara h 2 sIgE (kU/L) |
|---|---|---|---|---|---|---|
| **H7** | F | 8 | Allergic | 3.162 | >100 | >100 |
| **H8** | F | 13 | Allergic | 1.852 | >100 | 95 |
| **H10** | F | 12 | Allergic | 3.890 | >100 | >100 |
| **H11** | F | 14 | Allergic | 1.283 | >100 | >100 |
| **H13** | F | 10 | Allergic | 5.964 | >100 | >100 |
| **NA8** | F | 10 | Non-allergic | 0.360 | n.a. | n.a. |
| **NA9** | F | 14 | Non-allergic | 0.302 | n.a. | n.a. |
| **NA10** | F | 14 | Non-allergic | 0.167 | n.a. | n.a. |

**H**: PA children with PsIgE>100, **NA**: Non-allergic children
n.a.: not available

**Table S3**.
Top 10 enriched gene ontologies in cluster 5.

| Term | Ontology | Overlap | Adj. P | Genes |
|---|---|---|---|---|
| cytokine-mediated signaling pathway | GO:0019221 | 24/621 | 6.20E-06 | CD86 ITGB1 IL4R ANXA2 TRADD EBI3 ITGB2 MIF PSMA7 PSMB8 RBX1 PSMB9 PYCARD FCER2 AIM2 TMSB4X UBE2N HLA-DPB1 CD27 PTPN6 LTB PLP2 HLA-DQA1 HLA-DPA1 |
| antigen receptor-mediated signaling pathway | GO:0050851 | 11/185 | 0.00071 | BLK LAT2 MEF2C PDE4D UBE2N HLA-DPB1 HLA-DQA1 PSMA7 PSMB8 HLA-DPA1 PSMB9 |
| tumor necrosis factor-mediated signaling pathway | GO:0033209 | 9/116 | 0.00071 | PYCARD AIM2 TRADD TMSB4X CD27 LTB PSMA7 PSMB8 PSMB9 |
| regulation of T cell activation | GO:0050863 | 6/47 | 0.0018 | PYCARD LGALS3 PRELID1 LAPTM5 HLA-DPB1 HLA-DPA1 |
| neutrophil degranulation | GO:0043312 | 16/481 | 0.0026 | CD53 CAP1 CR1 ANXA2 HVCN1 TUBB ITGB2 MIF PYCARD VAMP8 LGALS3 RAB31 BIN2 SELL PTPN6 S100A11 |
| neutrophil activation involved in immune response | GO:0002283 | 16/485 | 0.0026 | CD53 CAP1 CR1 ANXA2 HVCN1 TUBB ITGB2 MIF PYCARD VAMP8 LGALS3 RAB31 BIN2 SELL PTPN6 S100A11 |
| neutrophil mediated immunity | GO:0002446 | 16/488 | 0.0026 | CD53 CAP1 CR1 ANXA2 HVCN1 TUBB ITGB2 MIF PYCARD VAMP8 LGALS3 RAB31 BIN2 SELL PTPN6 S100A11 |

| | | | | |
|---|---|---|---|---|
| mitochondrial ATP synthesis coupled proton transport | GO:0042776 | 4/17 | 0.0026 | ATP5PF ATP5PB ATP5MC3 ATP5F1E |
| regulation of interferon-gamma production | GO:0032649 | 7/86 | 0.0026 | PYCARD CR1 PDE4D EBI3 LAPTM5 HLA-DPB1 HLA-DPA1 |
| ATP synthesis coupled proton transport | GO:0015986 | 4/19 | 0.0038 | ATP5PF ATP5PB ATP5MC3 ATP5F1E |

**Table S4.**
Top 10 significantly enriched gene ontologies of PA subjects in cluster 5, compared to non-PA subjects in cluster 5.

| Term | Ontology | Overlap | Adj. P | Genes |
|---|---|---|---|---|
| cytokine-mediated signaling pathway | GO:0019221 | 32/621 | 2.70E-09 | SP100 SPI1 CDKN1B RPLP0 ITGB2 TNFRSF13C HIF1A RELA CNN2 SOCS1 HNRNPDL ALOX5 UBC PIM1 B2M JUNB JAK1 HSPA8 HLA-DRB5 HLA-B HLA-C IL16 HLA-A YWHAZ HLA-E HNRNPF IRF7 GRB2 PTPN6 LTB P4HB HLA-DQB1 |
| neutrophil degranulation | GO:0043312 | 27/481 | 8.80E-09 | VCP HSP90AB1 SLC44A2 ITGB2 CNN2 CYB5R3 LAMP1 FTH1 ALOX5 PSAP TBC1D10C COTL1 B2M HSPA8 DBNL JUP HLA-B HLA-C CYBA TMC6 EEF2 CKAP4 EEF1A1 PFKL PTPRC PKM PTPN6 |
| neutrophil activation involved in immune response | GO:0002283 | 27/485 | 8.80E-09 | VCP HSP90AB1 SLC44A2 ITGB2 CNN2 CYB5R3 LAMP1 FTH1 ALOX5 PSAP TBC1D10C COTL1 B2M HSPA8 DBNL JUP HLA-B HLA-C CYBA TMC6 EEF2 CKAP4 EEF1A1 PFKL PTPRC PKM PTPN6 |
| neutrophil mediated immunity | GO:0002446 | 27/488 | 8.80E-09 | VCP HSP90AB1 SLC44A2 ITGB2 CNN2 CYB5R3 LAMP1 FTH1 ALOX5 PSAP TBC1D10C COTL1 B2M HSPA8 DBNL JUP HLA-B HLA-C CYBA TMC6 EEF2 CKAP4 EEF1A1 PFKL PTPRC PKM PTPN6 |
| interferon-gamma-mediated signaling pathway | GO:0060333 | 10/68 | 2.30E-06 | SP100 HLA-DRB5 HLA-B IRF7 HLA-C HLA-A B2M JAK1 HLA-E HLA-DQB1 |
| antigen processing and presentation of exogenous peptide antigen via MHC class I, TAP-independent | GO:0002480 | 5/8 | 3.60E-06 | HLA-B HLA-C HLA-A B2M HLA-E |
| cellular response to cytokine stimulus | GO:0071345 | 22/482 | 1.40E-05 | RIPOR2 HSPA8 CDKN1B DUSP1 ITGB2 CXCR4 IL16 HIF1A YWHAZ RELA ZFP36L2 ZFP36L1 SIGIRR ZFP36 SOCS1 ALOX5 UBC PIM1 GRB2 PTPN6 JUNB JAK1 |

| | | | | |
|---|---|---|---|---|
| regulation of mRNA stability | GO:0043488 | 12/146 | 3.40E-05 | HSPA8 ZFP36 FUS YWHAB THRAP3 UBC CIRBP YBX1 PABPC1 YWHAZ ZFP36L2 ZFP36L1 |
| ribonucleoprotein complex assembly | GO:0022618 | 11/136 | 0.00011 | SETX RPL5 EIF2S3 HSP90AB1 SF3A2 PTGES3 RPLP0 EIF4H RPSA NOP53 EIF4B |
| antigen processing and presentation of endogenous peptide antigen via MHC class I via ER pathway | GO:0002484 | 4/7 | 0.00011 | HLA-B;HLA-C;HLA-A;HLA-E |

**Table S5.**
Demographics of samples used for Ara h 2 and DT binder sorting.

| Sample ID | Sex | Age | Allergic status | total IgE (ug/ml) | Peanut sIgE (kU/L) | Ara h 2 sIgE (kU/L) |
|---|---|---|---|---|---|---|
| **H10** | F | 12 | Allergic | 3.890 | >100 | >100 |
| **H11** | F | 14 | Allergic | 1.283 | >100 | >100 |
| **H16** | M | 10 | Allergic | 4.968 | >100 | >100 |
| **H17** | M | 11 | Allergic | 1.264 | >100 | 45 |
| **H18** | F | 11 | Allergic | 15.167 | >100 | >100 |
| **H19** | F | 23 | Allergic | 1.567 | >100 | 75.1 |
| **H20** | F | 23 | Allergic | 1.546 | >100 | 69.5 |
| **H22** | M | 17 | Allergic | 5.186 | >100 | >100 |
| **L7** | M | 14 | Allergic | 0.756 | 3.79 | 0.55 |
| **L8** | M | 8 | Allergic | 0.182 | 4.27 | 1.4 |
| **L9** | F | 12 | Allergic | 0.007 | 3.09 | 3.35 |
| **L17** | M | 17 | Allergic | 0.940 | 4.65 | <0.1 |
| **L18** | F | 30 | Allergic | 0.009 | 0.92 | 0.48 |
| **NA6** | F | 19 | Non-allergic | 0.021 | n.a. | n.a. |
| **NA15** | M | 18 | Non-allergic | 0.055 | n.a. | n.a. |

**H**: PA children with PsIgE>100, **L**: PA children with PsIgE<5, **NA**: Non-allergic children
n.a.: not available

**Table S6.**

Oligonucleotides and immunoglobulin (Ig) sequences used in this study.

Template switch and cDNA amplification

| | |
|---|---|
| Template switching oligo (TSO) | 5'-GCTAATCATTGCAAGCAGTGGTATCAACGCAGAGTACAT rGrGrG-3' |
| TSO-Oligo-dT30VN | 5'-GCTAATCATTGCAAGCAGTGGTATCAACGCAGAGTACAT TTTTTTTTTTTTTTTTTTTTTTTTTTTTTTVN-3' |
| cDNA amplification primer | 5'-GCTAATCATTGCAAGCAGTGGTATC-3' |

1st PCR to amplify BCRs

| | |
|---|---|
| Forward | 5'-CATTGCAAGCAGTGGTATCAACG-3' |
| IgM outer | 5'-CATGACGTCCTTGGAAGGCA-3' |
| IgGs outer | 5'-TTGTCCACCTTGGTGTTGCT-3' |
| IgAs outer | 5'-CAGGGCACAGTCACATCCT-3' |
| IgE outer | 5'-GTCGCAGGACGACTGTAAG-3' |
| IgK outer | 5'-TTCTCGTAGTCTGCTTTGCTCAG-3' |
| IgL_outer | 5'-GTCAGGCTCAGGTAGCTGC-3' |

2nd PCR to amplify BCR with barcodes

| | |
|---|---|
| 5'_plate_Row_ID primer | TACACGACGCTCTTCCGATCT (PLATE-BARCODE) GA (Row-barcode) GCTAATCATTGCAAGCAGTGGTATCAAC |
| 3'_ColimnID_IgInner primer | CTGCTGAACCGCTCTTCCGATC (Column barcode) IgInner |

Ig Inner primers

| | |
|---|---|
| IgM inner | 5'-CACGCTGCTCGTATCCGA-3' |
| IgGs inner | 5'-TCCTGAGGACTGTAGGACAGC-3' |
| IgAs inner | 5'-GGGAAGTTTCTGGCGGTCA-3' |
| IgE inner | 5'-GTGTCCCAGGTCACCATCAC-3' |
| IgK inner | 5'-GCGTTATCCACCTTCCACTGT-3' |
| IgL inner | 5'-TAGCTGCTGGCCG-3' |

PLATE-BARCODEs

| | |
|---|---|
| P1 | GCAGA |
| P2 | TCGAA |
| P3 | AACAA |
| P4 | GGTGC |
| P5 | CATTC |
| P6 | CGGTT |
| P7 | ATCCT |
| P8 | ATGTC |

Row-barcodes

| | |
|---|---|
| A | TAAGC |
| B | TGCAC |
| C | CTCAG |
| D | GGAAT |

| | |
|---|---|
| E | CGAGG |
| F | AGGAG |
| G | TGTTG |
| H | CAACT |

Column barcodes

| | |
|---|---|
| 1 | GTTCA |
| 2 | CAGGA |
| 3 | GTATA |
| 4 | CCTGT |
| 5 | ACCGC |
| 6 | ACTTA |
| 7 | GCTAG |
| 8 | GACGT |
| 9 | GGCTA |
| 10 | GAATG |
| 11 | CCAAC |
| 12 | GAGAC |

**Table S7.**

Oligonucleotide sequences used for real-time PCR reactions.

| **Target** | | **Sequences (5′–3′)** |
|---|---|---|
| *β-actin* | Forward | CACCATTGGCAATGAGCGGTTC |
| *β-actin* | Reverse | AGGTCTTTGCGGATGTCCACGT |
| *CD23/FCER2* | Forward | CCAGGAATTGAACGAGAGGAAC |
| *CD23/FCER2* | Reverse | TTGATCCACTTTTCAGGGCAC |
| *GLT-IGHE* | Forward | ACCATCCACAGGCACCAAAT |
| *GLT-IGHE* | Reverse | AGGTGGCATTGGAGGGAATG |
| *Total-IGHE* | Forward | ATCAGCTTGCTGACCGTCTC |
| *Total-IGHE* | Reverse | TAAGATCTTCACGGTGGGCG |
